# Oncogenic RAS-driven α2 integrin induction under nutrient stress promotes cancer cell motility

**DOI:** 10.64898/2026.04.02.716145

**Authors:** Bian Yanes, Mona Nazemi, Zhe Bao, Rachele Bacchetti, Ifeoluwa Oyelade, Elena Rainero

## Abstract

Cancer metabolism rewiring is one of the hallmarks of cancer, enabling cancer cell survival in a nutrient deprived microenvironment. Key to this is nutrient scavenging where cancer cells rely on extracellular proteins, including extracellular matrix (ECM) components, to sustain their proliferation. ECM uptake is mediated by α2β1 integrin, however it is not clear how this process is controlled by nutrient availability. Here we demonstrated that amino acid starvation promoted ECM internalisation, by inducing the expression of α2 integrin. Mechanistically, starvation-driven RAS/MAPK pathway activation in cells harbouring oncogenic RAS mutations and mTOR inhibition increased α2 integrin, while the GCN2-depedent integrated stress response was not required. Functionally, elevated α2 integrin levels promoted cell adhesion and migration in nutrient starved cells. Finally, α2 integrin was found upregulated in pancreatic tumours and correlated with poor prognosis in pancreatic adenocarcinoma patients. Together, these data indicate that the nutrient-starved pancreatic cancer microenvironment synergises with KRAS mutation to drive pancreatic cancer aggressiveness.

## Introduction

During cancer progression, cancer cells are under stress caused by either extrinsic factors, such as nutrient availability, hypoxia or exposure to treatments, or intrinsic factors, including protein accumulation. Indeed, the tumour microenvironment (TME) in pancreatic tumours was found to be hypoxic and deprived of nutrients, glucose and amino acids (AAs)[1], while pancreatic cancer cells grown in Tumour Interstitial Fluid Media (TIFM), which recapitulates the intra-tumoral nutrient concentrations, presented an amino acid deprivation signature [2, 3]. This pushes cancer cells to adopt alternative strategies of nutrient acquisition, including nutrient scavenging [4]. Indeed, glutamine starvation was shown to promote mutated KRAS-dependent uptake of extracellular proteins, such as albumin, via macropinocytosis [5], while, albumin promoted the growth of KRAS^G12D^ pancreatic cancer cells lacking essential amino acids [1]. Furthermore, the internalisation of the extracellular matrix (ECM) components collagen I and collagen IV supported the survival of pancreatic cancer cells under glucose or glutamine starvation [6]. Consistently, we demonstrated that the ECM supported breast cancer cell growth under AA deprivation, through an ECM uptake-dependent mechanism [7]. However, it is not clear how nutrient starvation promotes ECM scavenging.

Integrins are major ECM receptors. They are heterodimers consisting of two subunits, α and β. So far, there are 18 α subunits and 8 β subunits identified in mammals, which can form at least 24 integrins [8]. Importantly, integrins have been shown to mediate the internalisation of ECM components [9–13]. We and others have shown that α2β1 integrin promotes collagen I internalisation in different cell types, including fibroblasts, breast, pancreatic and ovarian cancer cells [10, 14–16] Moreover glucose deprivation triggered the internalisation of α5β1 intergin bound to fibronectin in ovarian cancer cells [17]. Finally, serum starvation *in vitro* and dietary restriction *in vivo* were shown to promote β4 integrin-dependent laminin uptake [18]. While β4 and αvβ3 integrins were shown to be upregulated by serum and glucose starvation, respectively [18, 19], little is known about the impact of AA deprivation impact on integrin expression.

Here we show that AA starvation promoted α2β1 integrin-dependent collagen I uptake, resulting in increased breast and pancreatic cancer cell growth. This was underpinned by a strong upregulation of α2 integrin, both at the mRNA and protein levels. Mechanistically, we found that α2 integrin expression was promoted by the RAS-MEK-ERK pathway in cells harbouring RAS mutations and by mTOR inhibition, while the integrated stress response did not play a role. Consistently, we observed an increase in ERK phosphorylation under AA starvation. Furthermore, AA starvation promoted pancreatic cancer cells adhesion to collagen I and migration. Together, these data demonstrate that AA starvation potentiates mutant RAS signalling, promoting α2 integrin expression, which in turn facilitate ECM scavenging, cell proliferation and migration.

## Results

### Amino acid starvation promoted a2b1 integrin-dependent ECM uptake

We recently demonstrated that ECM internalisation promoted breast cancer cell growth under amino acid starvation, by increasing cell division [7]. To investigate whether reduced nutrient availability was driving ECM uptake, we starved MCF10CA1 breast (pink ribbon) and SW1990 pancreatic (purple ribbon) cancer cells in amino acid free media for 18hr and seeded them on fluorescently labelled collagen I in either complete or AA-free media. While we detected some collagen I-positive vesicles in cells in complete media, more vesicles accumulated in AA-starved cells. Indeed, the uptake of collagen I was significantly increased when cells were grown under AA starvation compared to complete media (Figure 1A,B).

**Figure 1.**
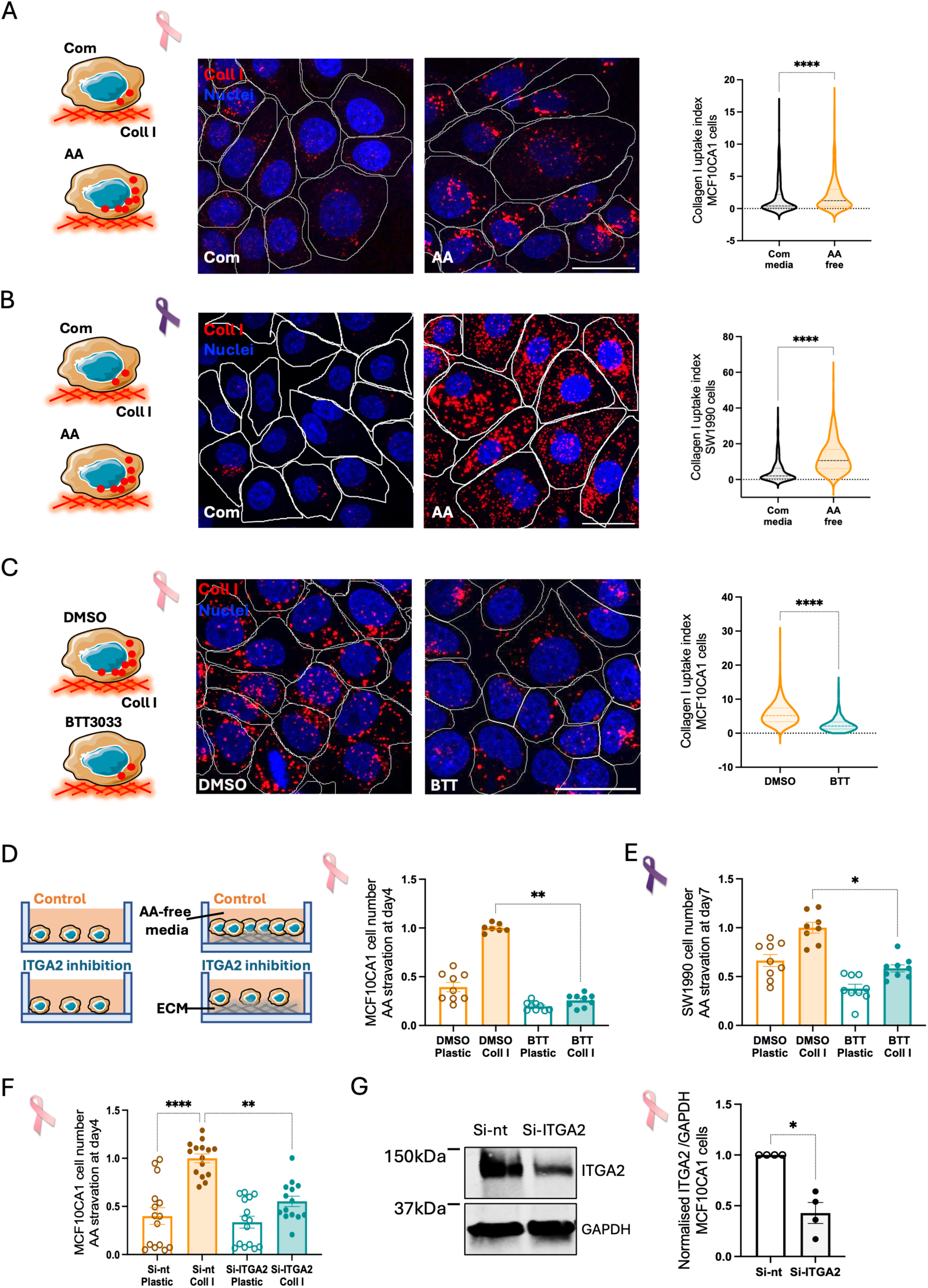
α**2 integrin was required for ECM uptake and ECM-dependent cell growth under amino acid starvation. (A,B)** MCF10CA1 (A) and SW1990 (B) cells were grown in complete media or amino acid-free media for 18hr and seeded on 1 mg/ml collagen I labelled with pH-rodo (red), in complete media (Com) or amino acid-free media (AA). Cells were stained with Hoechst 33342 (blue) and imaged live with a Nikon A1 confocal microscope, 60x magnification. White shapes highlight the cell edges. Scale bar 30 μm. Collagen I uptake index was calculated with Image J. N=3 independent experiments. ****p < 0.0001 Mann-Whitney test. **(C)** MCF10CA1 cells were grown in amino acid-free media (AA) for 18 hours and seeded in AA-free media for 6 hours on 1 mg/ml collagen I labelled with pH-rodo (red). DMSO or 15 μM BTT-3033 (BTT) were added 2 hours after seeding. Cells were stained with Hoechst 33342 (blue) and imaged live with a Nikon A1 confocal microscope, 60x magnification. White shapes highlight the cell edges. Scale bar 30 μm. Collagen I uptake index was calculated with Image J. N=3 independent experiments. ****p < 0.0001 Mann-Whitney test. **(D)** MCF10CA1 cells were seeded on 2 mg/ml collagen I in amino acid-free media (AA) for 4 days in the presence of DMSO (control) or 15 μM BTT-3033 (BTT). Cells were fixed, stained with Hoechst 33342 and imaged with an ImageXpress micro. Data were analysed with MetaXpress. N=3 independent experiments, graph shows mean ± SEM. **p < 0.01 Kruskal-Wallis, Dunn’s multiple comparisons test. **(E)** SW1990 cells were seeded on 2 mg/ml collagen I in amino acid-free media (AA) for 7 days in the presence of DMSO (control) or 10 μM BTT-3033. Cells were fixed, stained and imaged as in **D**. N=3 independent experiments, graph shows mean ± SEM. *p < 0.05 Kruskal-Wallis, Dunn’s multiple comparisons test. **(F)** MCF10CA1 cells were seeded on 2 mg/ml collagen I, transfected with an siRNA targeting α2 integrin (Si-ITGA2) or a non-targeting siRNA (Si-nt) as a control and grown for 4 days in amino acid-free media (AA). Cells were fixed, stained and imaged as in **D**. N=5 independent experiments, graph shows mean ± SEM. **p < 0.01, ****p < 0.0001 Kruskal-Wallis, Dunn’s multiple comparisons test. **(G)** MCF10CA1 cells were seeded on plastic, transfected with an siRNA targeting α2 integrin (si-ITGA2) or a non-targeting siRNA (si-nt) as a control. Cell lysates were collected after 72 hours, samples were run by Western Blotting, membranes were stained for α2 integrin (ITGA2) and GAPDH and imaged with a Licor Odyssey Sa system. The protein band intensity was measured with Image Studio Lite software. N=4 independent experiments, graph shows mean ± SEM. *p =0.0286 Mann-Whitney test. Pink and purple ribbons are from wikicommons, http://creativecommons.org/licenses/by-sa/3.0/.

We previously showed that α2β1 integrin was required for ECM internalisation in complete media, in both breast and pancreatic cancer cell lines [20]. To investigate if this was also the case under AA starvation, MCF10CA1 cells were seeded on fluorescently labelled collagen I in the presence or absence of BTT-3033, a pharmacological inhibitor preventing α2β1 integrin collagen binding, in AA free media (Figure 1C). In the presence of DMSO, several collagen I-containing endosomes were present inside the cells, while BTT-3033 treatment drastically reduced the amount of internalised collagen I. Indeed, image analysis demonstrated that α2β1 integrin pharmacological inhibition significantly reduced collagen I uptake under AA starvation.

To investigate whether the ECM-dependent cell growth under starvation was mediated by α2β1 integrin, MCF10CA1 cells were seeded on collagen I for 4 days under AA starvation (Figure 1D) in the presence or absence of BTT-3033. While collagen I promoted cell proliferation in the presence of DMSO, pharmacological inhibition of α2β1 integrin significantly prevented collagen I-dependent cell growth under starvation. Similarly, in SW1990 pancreatic cancer cells, collagen I-dependent cell growth was significantly impaired by α2β1 integrin inhibition (Figure 1E). To confirm the role of α2 integrin in ECM- dependent cell growth, MCF10CA1 cells were transfected with an siRNA targeting α2 integrin or a non- targeting siRNA control and seeded on collagen I for 4 days under AA starvation (Figure 1F). Similarly to the inhibitor results, α2 integrin knockdown significantly reduced collagen I-dependent cell growth under AA starvation. Western blot analysis confirmed a significant reduction of α2 protein levels in knockdown cells (Figure 1G).

Overall, these results indicate that ECM uptake is promoted by AA deprivation and ECM- dependent cell growth under starvation is mediated by α2β1 integrin.

### Amino acid starvation strongly increased a2 integrin expression at the mRNA and protein level

Some integrin heterodimers have been shown to be upregulated by nutrient starvation [18, 19], therefore we hypothesised that AA deprivation might promote collagen I uptake by stimulating α2β1 integrin expression. To investigate this, we performed qPCR in a panel of breast and pancreatic cancer cell lines to measure α2 integrin mRNA levels. Breast cancer cells (MCF10CA1, MDA-MB-231, and MCF7) and pancreatic cancer cells (PANC-1 and SW1990) were seeded in complete media or AA-free media for 18 hours. The mRNA was extracted and α2 integrin expression was measured (Figure 2A). The results show that AA starvation significantly boosted α2 integrin mRNA levels in all the cell lines. Indeed, we detected a 2-fold increase in MDA-MB-231 (Figure 2C) and PANC-1 cells (Figure 2E) and ∼3-fold increase in MCF10CA1 (Figure 2B), MCF7 (Figure 2D), and SW1990 cells (Figure 2F). The pancreatic cancer TME has been shown to be nutrient depleted and a reconstituted media which composition recapitulates the nutrient levels detected in the interstitial fluids of pancreatic tumours, TIFM, has been developed. Interestingly, RNAseq experiments detected an AA-starvation signature in cells grown in TIFM, compared to complete media [2, 3]. To determine if the reduced nutrient levels in TIFM were also promoting α2 integrin expression, SW1990 cells were seeded in complete media or TIFM for 18 hours and α2 integrin mRNA levels were measured by qPCR. Similarly to AA starvation, TIFM significantly increased α2 integrin expression (Figure 2G).

**Figure 2.**
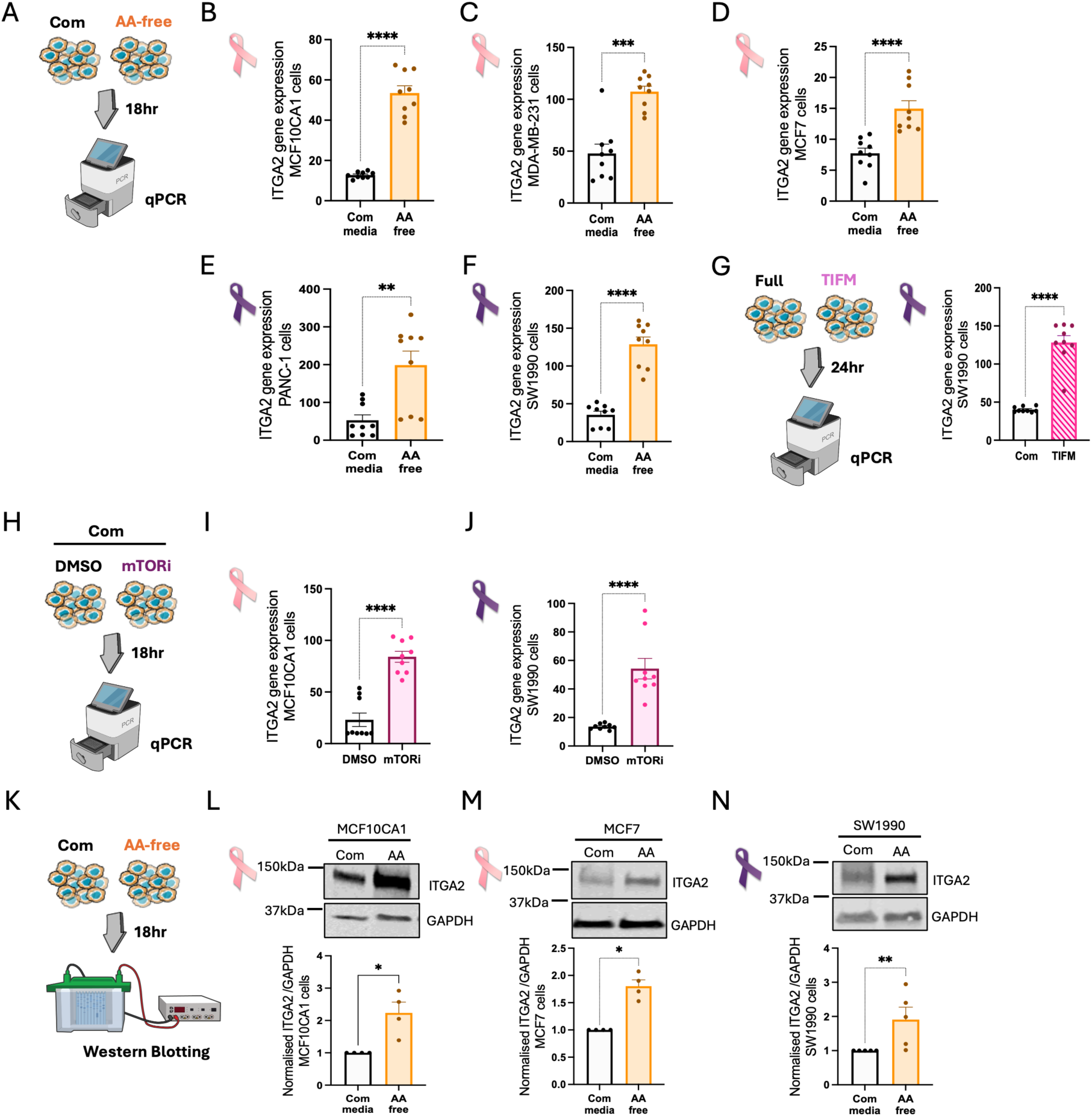
Amino acid starvation promoted. α**2 integrin expression. (A,G,H,K)** Schematic representation of experimental set up. Breast cancer cells **(B)** MCF10CA1, **(C)** MDA-MB-231, **(D)** MCF7 and pancreatic cancer cells **(E)** PANC-1 and **(F)** SW1990 were seeded in complete media (Com) or amino acid-free media (AA) for 18 hours. **(G)** SW1990 cells were seeded in complete media (Com) or TIFM for 18 hours. **(I)** MCF10CA1 and **(J)** SW1990 cells were seeded for 18 hours in complete media in the presence of DMSO as a control or 100 nM Torin 1 (mTORi). mRNA was extracted and α2 integrin (ITGA2) expression was measured by SYBR-Green qPCR. GAPDH was used as a control, and the data were plotted as 2^-ΔCT^. N=3 independent experiments, graphs show mean ± SEM. **p = 0.0040, ***p = 0.0005, ****p < 0.0001 Mann-Whitney test. **(L)** MCF10CA1, **(M)** MCF7 and **(N)** SW1990 cells were seeded in complete media (Com) or amino acid-free media (AA) for 18 hours. Cell lysates were collected, samples were run by Western Blotting, membranes were stained for α2 integrin (ITGA2) and GAPDH and imaged with a Licor Odyssey Sa system. The protein band intensity was quantified with Image Studio Lite software. N≥4 independent experiments, graphs show mean ± SEM. *p = 0.0286, **p = 0.0079 Mann-Whitney test. Pink and purple ribbons are from wikicommons, http://creativecommons.org/licenses/by-sa/3.0/; illustrations in A, G, H and K are with items from NIAID NIH BIOART, bioart.niaid.nih.gov/bioart/426 (A,G,H) and bioart.niaid.nih.gov/bioart/580 (K).

It is well established that AA deprivation deactivates the key nutrient sensor mechanistic target of rapamycin complex 1 (mTORC1), to promote cell anabolism [21]. Therefore, we wanted to investigate whether mTOR inhibition resulted in a similar upregulation of α2 mRNA levels. Cells were grown for 18 hours in complete media in the presence of the mTOR inhibitor Torin 1 or DMSO control (Figure 2H). Interestingly, Torin 1 significantly increased α2 integrin expression in both MCF10CA1 cells (Figure 2I) and SW1990 cells (Figure 2J).

To investigate whether AA starvation specifically controlled α2 integrin levels, we assessed the mRNA expression of other integrin subunits under AA starvation compared to complete media by qPCR (Figure S1A). We found that AA deprivation significantly induced the expression of β1 integrin in MCF10CA1 (Figure S1B), MDA-MB-231 (Figure S1C), and PANC-1 cells (Figure S1E), while in MCF7 (Figure S1D) and SW1990 cells (Figure S1F) there was a small, but not statistically significant, increase in β1 integrin mRNA levels.

Moreover, the expression of α3, α5 and α6 integrin did not change under AA starvation compared to complete media in MCF10CA1 cells (Figure S1G-I), while AA deprivation resulted in an increase in α5, but not α3 or α6, integrin expression in SW1990 cells (Figure S1J-L). These data indicate that AA starvation predominantly controlled α2 and, to a lesser extent, β1 integrin levels in both breast and pancreatic cancer cells.

To investigate whether the increase in α2 integrin mRNA was coupled with elevated protein expression, breast and pancreatic cancer cells were seeded in complete media or AA-free media for 18 hours and α2 integrin protein levels were measured by Western Blotting (Figure 2K). Consistent with our qPCR data, the expression of α2 integrin was significantly increased (∼ 2-fold) at the protein level under AA starvation in MCF10CA1 (Figure 2L), MCF7 (Figure 2M) and SW1990 cells (Figure 2N).

Overall, these results demonstrate that AA deprivation boosted the expression of α2 integrin at the mRNA and the protein levels, in both breast and pancreatic cancer cells, while α3, α5, and α6 integrin were not affected.

### Amino acid starvation promoted α2 integrin expression in 3D systems

3D culture represents a more physiological model [22]. Therefore, we wanted to investigate whether AA starvation upregulated α2 integrin expression in 3D systems as well. MCF10CA1 cells (Figure 3A) and SW1990 cells (Figure 3B) were grown in Matrigel for 24 hours in complete media, starved for 2 days in AA-free media and α2 integrin levels were measured by immunofluorescence. Filled acini structures appeared slightly irregular and smaller under AA starvation compared to complete media, with α2 integrin being localised at the cell surface. Image analysis indicated that α2 integrin signal intensity was significantly increased under AA starvation compared to complete media in both cell lines (Figure 3A,B). As expected, the acini area was reduced upon AA starvation (Figure 3A,B). These results indicate that AA deprivation increased α2 integrin expression also in 3D systems.

**Figure 3.**
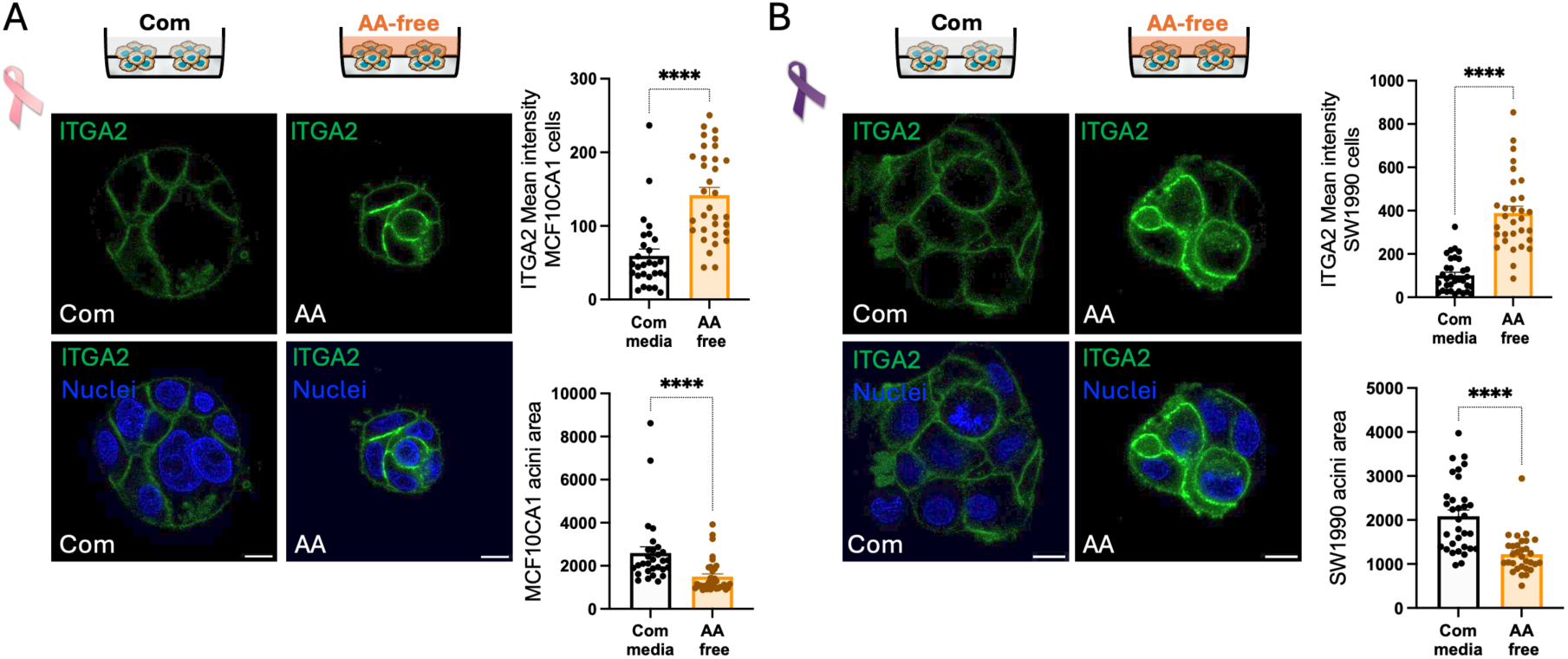
Amino acid starvation promoted. α**2 integrin expression in 3D. (A)** MCF10CA1 and **(B)** SW1990 cells were grown in 3D Matrigel. After 24 hours, media was changed to either complete (Com) or amino acid-free media (AA). Cells were incubated for 2 days, fixed and stained for α2 integrin (ITGA2, green) and nuclei (blue). Cells were imaged with a Nikon A1 confocal microscope, 60x objective. Scale bar 10 μm. ITGA2 mean intensity and acini area were measured using Image J. N=3 independent experiments, graphs show the mean ± SEM. ****p < 0.0001 Mann-Whitney test. Pink and purple ribbons are from wikicommons, http://creativecommons.org/licenses/by-sa/3.0/.

### Amino acid starvation promoted a2 integrin expression in a RAS-dependent and GCN2- independent manner

AA starvation promotes the activation of the general control nonderepressible 2 (GCN2)- mediated integrated stress response, leading to changes in gene expression underpinning metabolic adaptations to nutrient deprivation [23] (Figure 4A). Therefore, we reasoned that GCN2 might be responsible for α2 integrin expression induced by the lack of AAs. To confirm that GCN2 was activated by AA starvation in SW1990 cells, we measured the expression of asparagine synthetase (ASNS), a well-established GCN2 target gene [24], by qPCR. As expected, the expression of ASNS was significantly increased under AA starvation compared to complete media (Figure S2A). To confirm the involvement of GCN2, SW1990 cells were seeded under AA starvation for 18 hours in the presence or absence of the GCN2 inhibitor GCN2iB. As expected, the starvation-induced increase in ASNS expression was completely abolished by GCN2 inhibition (Figure S2B). To investigate whether the starvation-induced increase in α2 integrin expression was mediated by GCN2, breast and pancreatic cancer cells were seeded in AA-free media for 18 hours in the presence or absence of the GCN2 inhibitor and α2 integrin mRNA level was measured by qPCR (Figure 4B). Interestingly, the inhibition of GCN2 resulted in small but statistically significant reduction in the expression of α2 integrin in MCF7 (Figure 4E) but not in MCF10CA1 (Figure 4C) or MDA-MB-231 breast cancer cells (Figure 4D). Similarly, the inhibition of GCN2 did not affect α2 expression in both SW1990 (Figure 4H) and PANC-1 (Figure 4I) pancreatic cancer cells. The data indicate that the integrated stress response is not the main pathway promoting α2 integrin expression under AA starvation.

**Figure 4.**
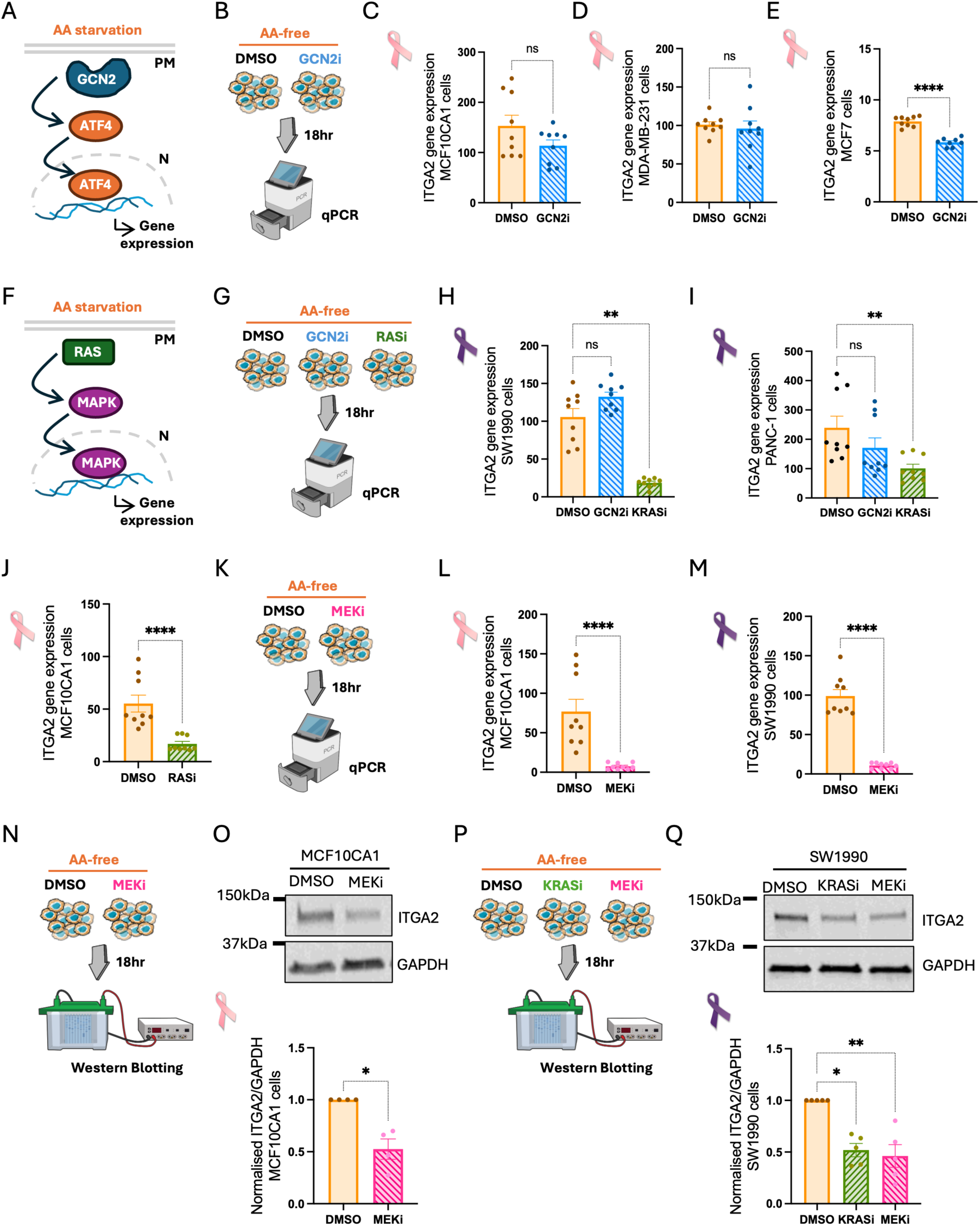
RAS/MAPK signalling, but not GCN2, was required for amino acid starvation-induced α2 integrin expression. (A,F) GCN2 and RAS signalling pathway schematics. (B,G,K,N,P) Schematic representation of experimental set up. (C-E) Breast cancer cells (C) MCF10CA1, (D) MDA-MB-231 and (E) MCF7 were grown in amino acid-free media (AA) for 18 hours in the presence of 5 μM GCN2iB (GCN2i) or DMSO as a control. mRNA was extracted and α2 integrin (ITGA2) was measured by SYBR-Green qPCR. GAPDH was used as a control, and the data were plotted as 2^-ΔCT^. N=3 independent experiments. Graphs show mean ± SEM, ns: Non-significant, ****p < 0.0001 Mann-Whitney test. (H,I) Pancreatic cancer cells (H) SW1990 and (I) PANC-1 were seeded in AA-free media for 18 hours in the presence of 5 μM GCN2iB (GCN2i), 200 nM MRTX1133 (KRASi) or DMSO as a control. (J) MCF10CA1 breast cancer cells were grown in AA-free media for 18 hours in the presence of 1 μM RMC-6236 (RASi) or DMSO as a control. (L) MCF10CA1 and (M) SW1990 cells were seeded in AA-free media for 18 hours in the presence of 10 μM Selumetinib (MEKi) or DMSO as a control. mRNA was extracted and α2 integrin (ITGA2) expression was measured by SYBR-Green qPCR. GAPDH was used as a control and the data were plotted as 2^-ΔCT^. N=3 independent experiments, graphs show mean ± SEM. **p < 0.01 Kruskal Wallis test (H, I); ****p < 0.0001 Mann-Whitney test (J, L, M). (O) MCF10CA1 and (Q) SW1990 cells were seeded in AA-free media in the presence or absence of 10 μM Selumetinib (MEKi), 200 nM MRTX1133 (KRASi) or DMSO as a control. Cell lysates were collected, and samples were run by Western Blotting. Membranes were stained for α2 integrin (ITGA2) and GAPDH and imaged with a Licor Odyssey Sa system. The protein band intensity was quantified with Image Studio Lite software. N≥4 independent experiments, graphs show the mean ± SEM, *p = 0.0286 Mann-Whitney test (N), *p < 0.05, **p < 0.01 Kruskal Wallis multiple comparison test (P). Pink and purple ribbons are from wikicommons, http://creativecommons.org/licenses/by-sa/3.0/; illustrations in B,G,K,N, and P are with items from NIAID NIH BIOART, bioart.niaid.nih.gov/bioart/426 (B,G,K) and bioart.niaid.nih.gov/bioart/580 (N,P).

RAS is an oncogene mutated in multiple cancer types [25]. RAS cycles between an inactive GDP- bound state and an active GTP-bound state. Most RAS mutations occur at position G12, G13, and Q61, blocking RAS ability to hydrolyse GTP to GDP and therefore preventing inactivation [26]. All the cells used in this study harbour a mutation in one of the RAS isoforms, apart from MCF7 cells. MCF10CA1 cells are HRAS^G12V^, MDA-MB-231 cells are KRAS^G13D^, and both SW1990 and PANC-1 cells are KRAS^G12D^. Interestingly, mitogen activated protein kinase (MAPK) activation downstream of RAS has been reported to regulate the expression of GCN2-independent genes under AA starvation [27] (Figure 4F). To test whether the increase in α2 integrin expression was driven by RAS, PANC-1 and SW1990 cells were seeded in AA-free media for 18 hours in the presence or absence of a newly developed KRAS^G12D^ inhibitor, MRTX1133 [28], while MCF10CA1 cells were seeded in the presence or absence of RMC-6236, a Pan-RAS inhibitor [29] (Figure 4G) and α2 integrin mRNA levels were measured. RAS pharmacological inhibition significantly reduced the expression of α2 integrin in SW1990 (Figure 4H), PANC-1 (Figure 4I) and MCF10CA1 cells (Figure 4J). To determine whether the role of RAS was AA starvation-specific, pancreatic cancer cells were treated with MRTX1133 in complete media for 18 hours (Figure S3A). We found that KRAS inhibition markedly reduced α2 integrin expression in SW1990 cells (Figure S3B) but not in PANC-1 cells (Figure S3C), while GCN2 inhibition did not affect α2 levels. The discrepancy in MRTX1133 response could be due to the fact that SW1990 cells are homozygous for KRAS^G12D^, while PANC1 cells are heterozygous. Therefore, the WT KRAS allele, which is not affected by MRTX1133, might sustain α2 integrin expression in complete media. Together, these data demonstrate that AA starvation promotes α2 integrin expression through a RAS-dependent signalling pathway.

### α2 integrin expression was regulated by MEK under amino acid starvation

RAS activation triggers different signalling pathways, including the RAS-RAF-MEK-ERK pathway. It was previously reported that extracellular signal-regulated kinase (ERK) can be activated by AA starvation [30]. Hence, to verify that this was the case in our system, we measured the phosphorylation of ERK upon AA starvation. MCF10CA1, SW1990 and PANC-1 cells were seeded in serum-free media for 18 hours. Cells were incubated in complete or AA-free media (without serum) for 5, 10, and 15 minutes and ERK and p-ERK protein levels were measured by Western Blotting. Consistently with previous observations, ERK phosphorylation increased significantly in MCF10CA1 (Figure S4A), SW1990 (Figure S4B) and PANC-1 cells (Figure S4C) within 5 min of starvation compared to complete media and then it returned to control levels. There was no difference in ERK protein levels between complete and AA-free media. This indicated that AA starvation promotes RAS/ERK signalling activation in breast and pancreatic cancer cells.

We then wanted to investigate whether the mitogen-activate protein kinase kinase (MEK)/ERK pathway was required for AA starvation-induced α2 integrin expression. MCF10CA1 breast cancer cells and SW1990 pancreatic cancer cells were seeded in AA-free media for 18 hours in the presence or absence of a MEK inhibitor, Selumetinib [31], and α2 integrin mRNA and protein levels were measured (Figure 4K,N,P). MEK inhibition profoundly reduced α2 integrin mRNA in both MCF10CA1 (Figure 4L) and SW1990 cells (Figure 4M). Similarly, MEK inhibition significantly reduced α2 integrin expression at the protein level in MCF10CA1 cells (Figure 4O), while both MEK and KRAS pharmacological inhibitors significantly reduced α2 protein levels in SW1990 cells (Figure 4Q).

Consistently, MEK inhibition in MCF10CA1 cells (Figure 5A) and KRAS inhibition in SW1990 cells (Figure 5B) strongly reduced the expression of α2 integrin in 3D systems. As expected, MRTX1133 significantly reduced the phosphorylation of ERK in both SW1990 cells (Figure S5A,B) and PANC-1 cells (Figure S5C) while Selumetinib completely abolished ERK phosphorylation in MCF10CA1 (Figure S5D,E), SW1990 (Figure S5F) and PANC-1 cells (Figure S5G).

**Figure 5.**
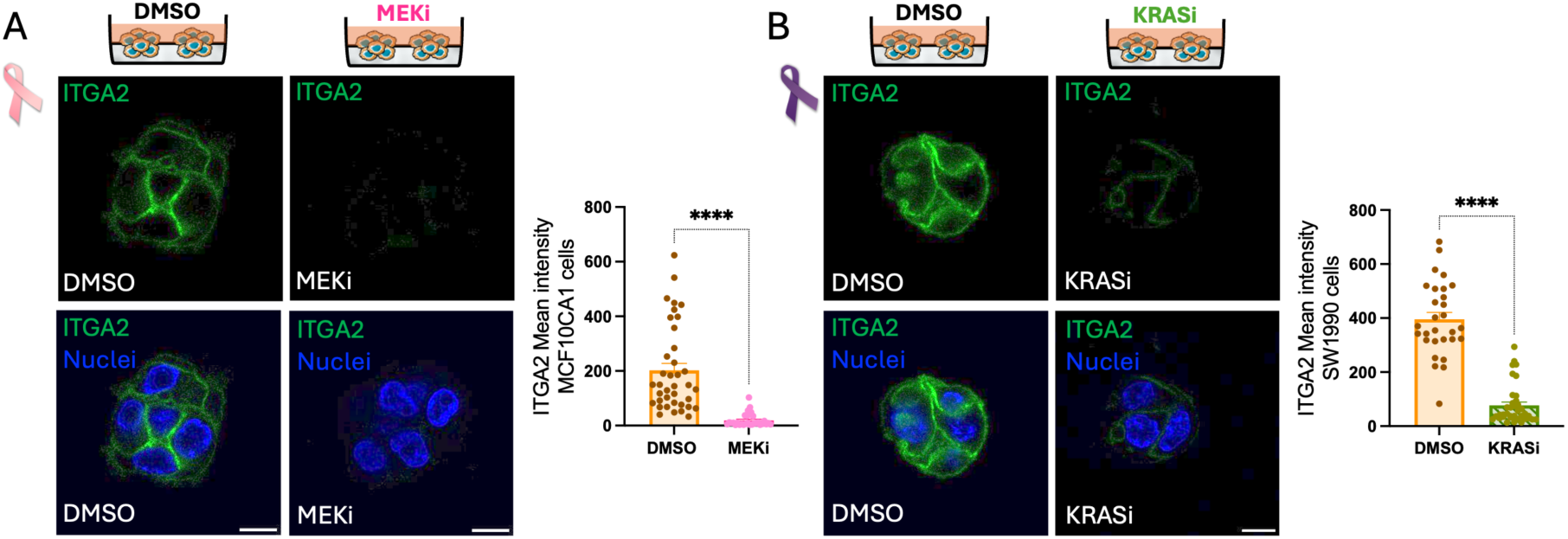
KRAS/MEK signalling was required for amino acid starvation-induced. α**2 integrin expression in 3D**. **(A)** MCF10CA1 cells and **(B)** SW1990 cells were grown in 3D Matrigel. After 24 hours, media was changed to amino acid free-media in the presence of 10 μM Selumetinib (MEKi), 200 nM MRTX1133 (KRASi) or DMSO control. Cells were incubated for 2 days, fixed and stained for α2 integrin (ITGA2, green) and nuclei (blue). Cells were imaged with a Nikon A1 confocal microscope, 60x objective. Scale bar 10 μm. ITGA2 mean intensity was measured using Image J. N=3 independent experiments, graphs show the mean ± SEM, ****p < 0.0001 Mann-Whitney test. Pink and purple ribbons are from wikicommons, http://creativecommons.org/licenses/by-sa/3.0/

Overall, these data demonstrate that α2 integrin expression is triggered by the AA starvation- dependent activation of the MEK/ERK signalling pathway downstream mutant RAS in breast and pancreatic cancer cells.

### Amino acid starvation promoted pancreatic cancer cell adhesion and migration

Integrins are cell surface receptors which promote cell adhesion and motility [32]. As the expression of the collagen I receptor α2 integrin, but not the fibronectin receptor α5 integrin or the laminin receptors α3 and α6 integrins, was increased under AA starvation, we hypothesised that AA starvation promoted cell adhesion to collagen I, but not to other substrates. To address this, SW1990 pancreatic cancer cells were grown in complete or AA-free media for 18 hours, then seeded on plastic, collagen I or Matrigel for 15, 30 or 60 minutes (Figure 6A). As expected, cell adhesion was increased over time on all substrates. However, we found that more AA-starved cells adhered to collagen I compared to cells in complete media within 30 minutes of adhesion (∼ 50% increase) (Figure 6C). Interestingly, the number of cells adhered to plastic or Matrigel was similar in both complete and AA free media (Figure 6B,D). These data indicate that AA starvation specifically promotes cell adhesion to α2 integrin substrates. To determine whether AA deprivation also promoted cell spreading, PANC-1 cells were grown in complete or AA free media for 18 hours, seeded on collagen I and imaged live (supplementary video 1 and 2). Still images from the videos show that AA-starved cells spread faster on collagen I compared to cells maintained in complete media (Figure S6).

**Figure 6.**
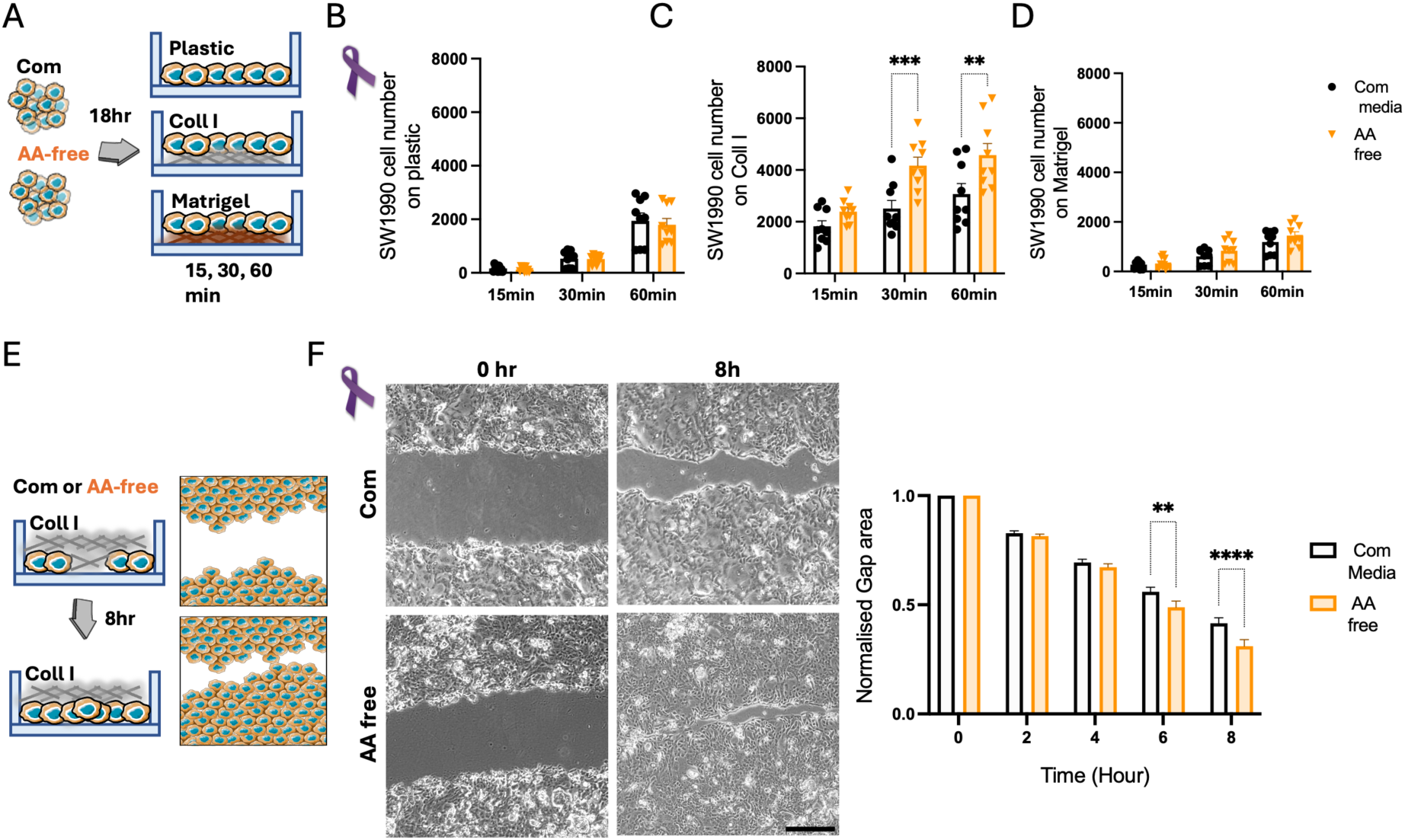
Amino acid starvation promoted cell adhesion to collagen I and cell migration. (A, **E)** Schematic representation of experimental set up. **(B-D)** SW1990 cells were grown in complete media (Com) or amino acid-free media (AA) for 18 hours then seeded on (B) plastic, (C) 2 mg/ ml collagen I (Coll I), or (D) 3 mg/ ml Matrigel. Cells were fixed after 15, 30, or 60 minutes and stained with Hoechst 33342. The cells were imaged using an Image Xpress micro system and cell numbers were quantified with MetaXpress and CME software. Graphs show mean ± SEM, N=3 independent experiments. **p < 0.01, ***p < 0.001 Two Way ANOVA, Tukey’s test. **(F)** SW1990 cells were seeded and incubated overnight to achieve confluency. The monolayer was scratched manually and 0.5 mg/ml collagen I in complete media (Com) or amino acid-free media (AA) was added to the scratched monolayer. The scratches were imaged after 0, 2, 4, 6, and 8 hours using an Olympus Inverted Fluorescence Microscope, 4x objective. Scale bar 250 μm. Area of the scratches was manually calculated by tracing the perimeter of the non-invaded area using Image J. Graph shows mean ± SEM, N=3 independent experiments. **p < 0.01, ****p < 0.0001 Two Way ANOVA, Tukey’s test.

Nutrient starvation has been suggested to promote cancer cell migration and metastasis [33, 34] and we demonstrated that α2β1 integrin-dependent ECM uptake was required for cell migration [20]. Therefore, we wanted to investigate whether AA starvation promoted pancreatic cancer cell migration. SW1990 cell monolayers were scratched, overlayed with collagen I and allowed to migrate in complete or AA-free media for up to 8 hours (Figure 6E). We observed that AA starvation enhanced SW1990 cells migration, as after 8 hours the scratches appeared fully closed in AA depleted media but not in complete media. Quantification of the scratched area indicated a significant reduction in AA- starved cells at both 6 and 8 hours, indicating increased cell migration (Figure 6F).

Overall, these data demonstrate that AA starvation promoted adhesion, spreading and migration in the presence of collagen I, the main α2 integrin substrate.

### Amino acid starvation promoted cell adhesion and ECM uptake in a MAPK-dependent manner

We hypothesised that the AA starvation-induced adhesion to collagen I and collagen I uptake were due to α2 integrin upregulation, which was dependent on RAS/MEK signalling. Therefore, we expected that RAS/MEK inhibition during the AA starvation period reduced collagen I adhesion and uptake. To test this, SW1990 pancreatic cancer cells were grown in AA-free media for 18 hours in the presence or absence of either Selumetinib (MEKi), MRTX1133 (KRASi) or DMSO control, then seeded on collagen I for 30 minutes in the absence of inhibitors. Consistently with our hypothesis, MEK and RAS inhibition resulted in a 25% reduction in cell adhesion to collagen I (Figure 7A).

**Figure 7.**
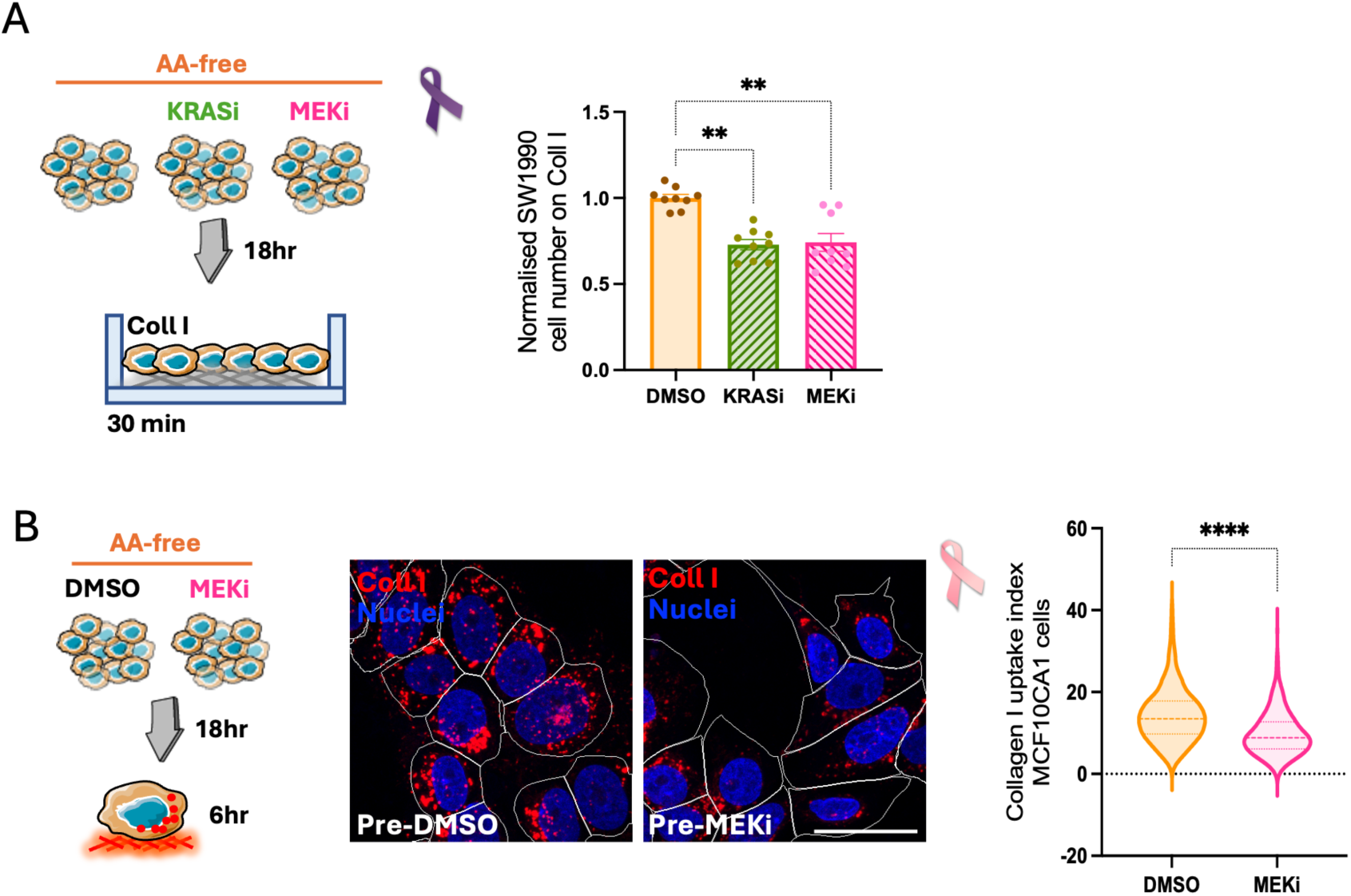
RAS/MAPK signalling was required for amino acid starvation-induced cell adhesion and collagen I uptake. **(A)** SW1990 cells were grown in amino acid-free media (AA) in the presence of 200 nM MRTX1133 (KRASi), 10 μM Selumetinib (MEKi) or DMSO for 18 hours and seeded on 2 mg/ml collagen I (Coll I) under AA starvation but in the absence of inhibitors. Cells were washed twice with PBS, fixed and stained with Hoechst 33342 after 30 minutes. Cells were imaged using an INCell Analyser 2200 and cell numbers were quantified with Cell Profiler. Graph shows mean ± SEM. N=3 independent experiments, **p < 0.001, ***p < 0.001 Kruskal Wallis test. **(B)** MCF10CA1 cells were grown in AA-free media in the presence of DMSO or 10 μM Selumetinib (MEKi) for 18 hours and seeded in AA-free media on pH-rodo labelled 1 mg/ml collagen I (Coll I, red) for 6 hours (in the absence of inhibitors). Cells were stained with Hoechst 33342 (blue) and were imaged live with a Nikon A1 confocal microscope, 60x magnification. White shapes highlight the cell edges. Scale bar 30 μm. Collagen I uptake index was calculated with Image J. Graph shows mean ± SEM. N=3 independent experiments, ****p < 0.0001, Mann-Whitney test. Pink and purple ribbons are from wikicommons, http://creativecommons.org/licenses/by-sa/3.0/

Similarly, MCF10CA1 breast cancer cells were grown in AA-free media for 18 hours in the presence or absence of Selumetinib (MEKi) and seeded on fluorescently labelled collagen I for 6 hours in the absence of the inhibitor. In DMSO-pretreated cells, several collagen I-containing endosomes can be seen inside the cells, while in MEKi-pretreated cells there were fewer collagen I-containing vesicles. Quantification of the collagen I uptake index showed that MEK inhibition during the AA starvation phase significantly reduced collagen I uptake under AA starvation (Figure 7B).

Overall, our data show that both cell adhesion and collagen I uptake under AA starvation were mediated by MEK-dependent α2 integrin upregulation.

### Integrin a2 was upregulated in pancreatic cancer and correlated with poor prognosis

Our data showed that α2 integrin expression was increased by AA deprivation in cancer cells. The TME in pancreatic cancer is known for being deprived of nutrients, including AAs [1]. Therefore, we assessed the expression of α2 integrin in pancreatic cancer patients. Interestingly, α2 integrin expression was highly increased in pancreatic adenocarcinoma tumours compared to healthy pancreatic tissue (Figure 8A). Moreover, high α2 expression correlated with a reduction in the patients’ overall survival (Figure 8B) and disease-free survival (Figure 8C), indicating that α2 integrin may contribute to tumour progression and survival in pancreatic cancer patients.

**Figure 8.**
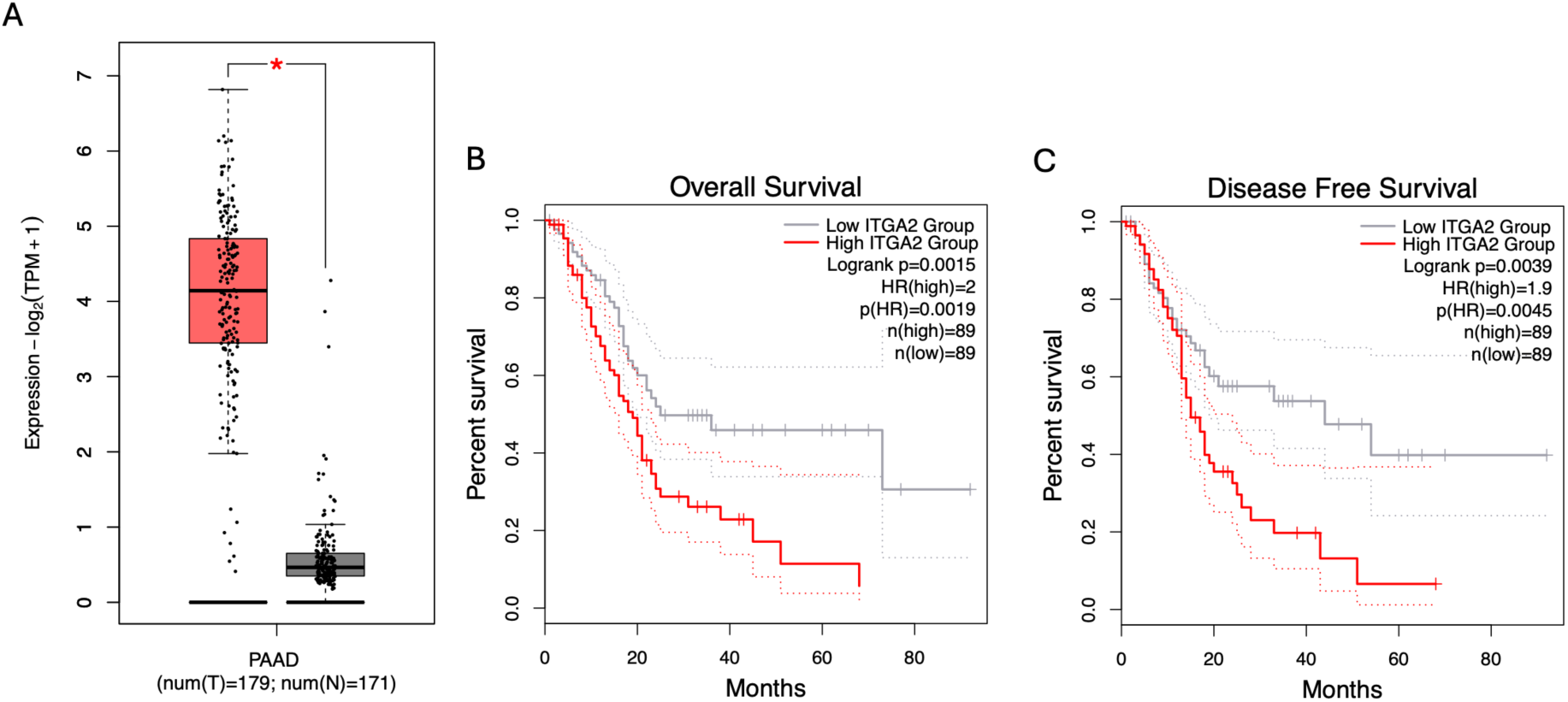
α2 integrin was overexpressed in pancreatic tumours and correlated with poor prognosis. **(A)** RNA sequencing data from pancreatic adenocarcinoma tumours (red, n=179) and normal pancreas (grey, n=171) for α2 integrin expression. **(B-C)** 178 pancreatic adenocarcinoma patients were stratified into low and high α2 integrin (ITGA2) expressors based on median gene expression. The Kaplan-Meier analysis compared overall survival **(B)** and disease-free survival **(C)** of patients with tumours expressing high (red) and low (grey) α2 integrin levels. The data were generated using GEPIA2 (http://gepia2.cancer-pku.cn/#index).

## Discussion

Nutrient scavenging, including the uptake of ECM components, plays a key role in supporting the growth of nutrient starved cancer cells [7]. Here we showed that collagen I internalisation was promoted by AA starvation, in an α2β1 integrin dependent manner. We demonstrated that AA deprivation promoted the activation of the MEK/ERK signalling pathway in breast and pancreatic cancer cells harbouring RAS mutations, which in turn increased α2 integrin expression. This underpinned increased adhesion, collagen I internalisation and cell migration induced by the lack of AA. Consistent with the fact that pancreatic tumours are nutrient deprived, α2 integrin levels were significantly upregulated in tumours compared to healthy pancreas and correlated with poor prognosis (Figure 9).

**Figure 9.**
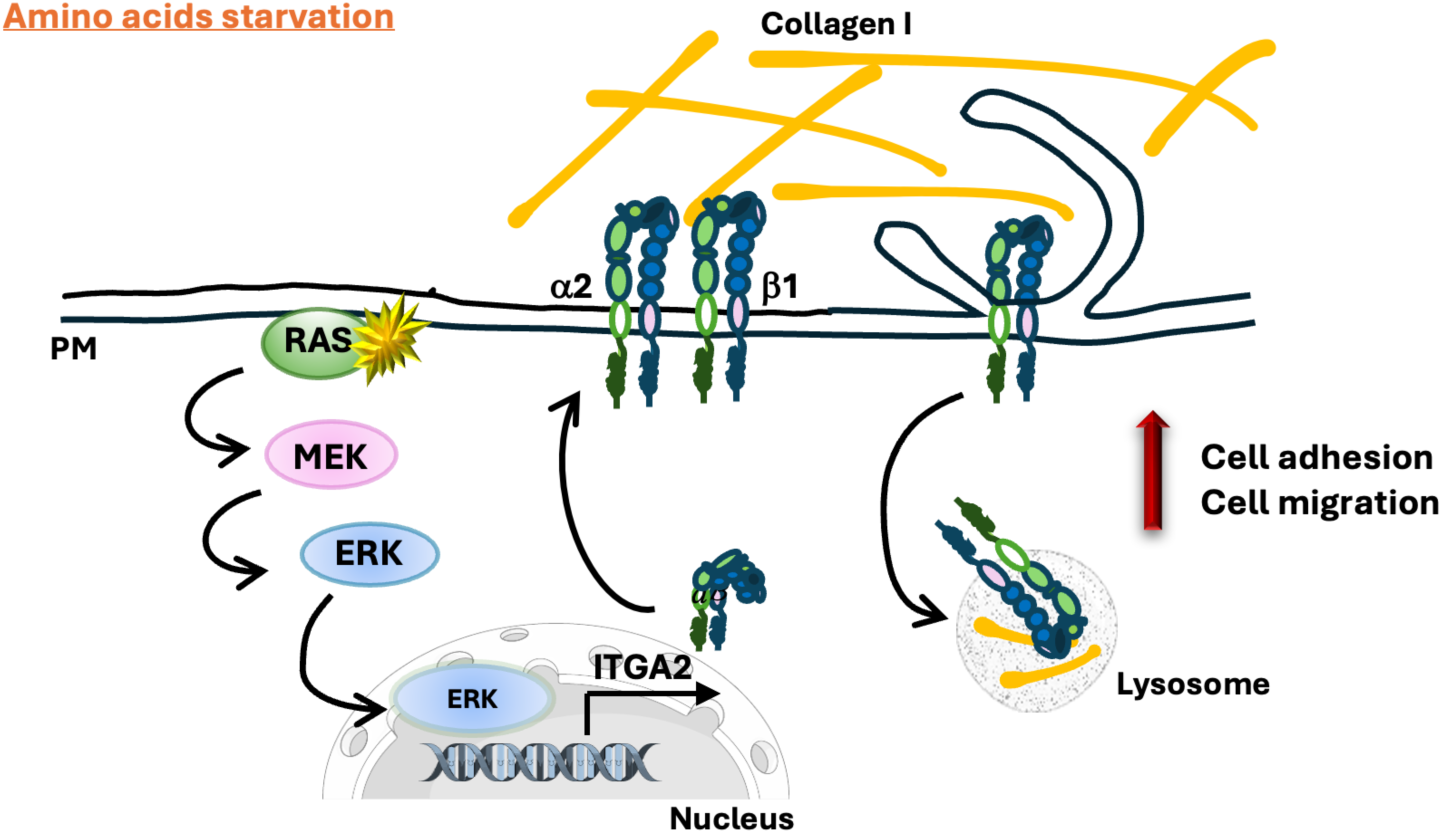
Working model. In breast cancer and pancreatic cancer cells harbouring RAS mutations, amino acid starvation induced the expression of α2 integrin (ITGA2), both at the mRNA and the protein level, via a RAS/MEK/ERK-dependent mechanism, therefore increasing α2β1 integrin binding to collagen I and its internalisation, which enhanced cell adhesion, collagen uptake and cell migration. PM, plasma membrane.

We found that collagen I internalisation was significantly induced by AA deprivation. Consistently, laminin endocytosis was shown to be promoted by serum starvation in mammary epithelial cells [18], glucose starvation increased fibronectin internalisation in ovarian cancer cells [17], while glucose and glutamine deprivation stimulated soluble collagen I and collagen IV uptake in pancreatic cancer cells [6]. This indicates that nutrient availability is a key determinant in stimulating ECM protein scavenging in different cell types.

We showed that collagen I uptake in breast cancer cells under AA starvation was mediated by α2β1 integrin. Consistently, we have recently demonstrated that α2β1 integrin controlled ECM internalisation in breast, pancreatic and ovarian cancer cells in complete media [20], indicating that similar mechanisms are likely to control ECM uptake regardless of nutrient availability. In agreement with the fact that ECM internalisation is required for ECM-dependent breast cancer cell growth under starvation [7], here we showed that α2β1 integrin was required for collagen I-dependent cell proliferation in both breast and pancreatic cancer cells under AA starvation.

α2β1 integrin is one of the main collagen binding receptors. The α2 subunit is responsible for binding to the ligand, and it determines the ligand specificity, while the β1 subunit indirectly binds to the cytoskeleton [35, 36]. Here, the expression of α2 integrin subunit was increased in a panel of breast and pancreatic cancer cells upon AA starvation, as well as in pancreatic cancer cells grown in TIFM, which more closely recapitulates intra-tumoral nutrient levels. β1 integrin was upregulated in some cell lines, while α3, α5 and α6 integrin levels did not change. The expression of different integrin isoform has been reported to be regulated by nutrient availability. Indeed, β4 integrin was increased in epithelial cells under serum starvation [18], while β3 integrin expression was promoted by glucose starvation in non-small cell lung cancer cells [19]. Moreover, low glucose promoted α2 and decreased αvβ3 integrin levels in human glomerular epithelial cells [37]. These observations support the fact that integrin expression is controlled by nutrient availability, however the molecular mechanisms behind this were not previously investigated.

It is well established that AA deprivation triggers the GCN2 branch of the integrated stress response [23]. Here we showed that the increase in α2 integrin expression under AA starvation was partially mediated by GCN2 only in wild-type RAS breast cancer cells (MCF7), while in both HRAS mutated (MCF10CA1) and KRAS mutated cells (PANC-1 and SW1990), α2 integrin expression was mediated by the RAS/MEK signalling pathway, in a GCN2-independent manner. Consistently, the upregulation of GCN2-independent genes has been reported in mouse embryonic fibroblasts under leucine starvation [24]. Moreover, mutant RAS has been shown to control integrin expression. For instance, the transformation of MDCK cells with HRAS^G12V^ or KRAS^G12V^ upregulated α6 integrin expression, through a MAPK/ERK/FOSL1-dependent pathway [38]. Similarly, pharmacological inhibition of MEK1 in pancreatic cancer cells and melanoma cells decreased the expression of α6 and β3 integrins. In addition, the sustained activation of the RAF-MEK-ERK pathway in NIH3T3 fibroblasts significantly increased α6 and β3 integrin levels, while transient ERK activation by growth factors did not affect β3 integrin expression [39]. Interestingly, the same study showed that the expression of α2 integrin was not increased by activating RAF-MEK-ERK pathway in fibroblasts [39], suggesting a cell type specific regulation. Our results also showed that the expression of α2 integrin in complete media was mediated by RAS only in SW1990, but not in PANC-1 cells. On the contrary, it has recently been demonstrated that the inhibition of either KRAS^G12D^, by the inhibitory peptide KRpep-2d, or ERK1/2 decreased α2 levels in PANC-1 and AsPC-1 cells [40]. This discrepancy could be due to the different inhibitors used. Moreover, Cai et al did not report the length on the inhibition, making it impossible to determine whether differences in the more or less prolonged RAS and ERK inhibition could have differential effects on α2 expression.

We showed that AA starvation transiently induced ERK phosphorylation and this is consistent with previous observations. Indeed, AA limitation promoted cFOS expression in HepG2 cells in a RAS/MAPK-dependent manner [27]. We hypothesise that this transient ERK activation is sufficient to drive the expression of immediate early transcription factors of the FOS, JUN and EGR families, which will then promote α2 integrin expression. Indeed, both FOS and EGR1 are predicted to bind to α2 promoter and were reported to be upregulate by AA starvation [41]. It is currently unknown how AA starvation triggers ERK activation. mTOR inhibition by Rapamycin has been shown to promote ERK phosphorylation in lung cancer cell lines [42]. Since we detected a similar increase in α2 integrin expression upon mTOR inhibition, it is possible that this is what triggers ERK activation. It would be important to further determine the crosstalk between mutant KRAS and mTOR signalling. Indeed, in multiple myeloma cells, oncogenic KRAS and NRAS have been reported to interact with the AA transporter SLC3A2 and mTOR, promoting mTORC1 activation [43]. Furthermore, AA or serum starvation has been shown to trigger AMPK activation in human erythroleukemia K562 cells and rat hepatoma H4IIE cells, which in turn activated MEK/ERK and induced autophagy [44]. However, the phosphorylation of ERK was shown at 6 hours of starvation, while we found that ERK phosphorylation was triggered after 5 minutes of AA starvation. This is in agreement with a previous study showing that starving fibroblasts in AA-free EBSS solution rapidly increased the phosphorylation of ERK after 5 minutes of starvation [30]. The authors also showed that the cell volume decreased upon starvation, suggesting that the physical shrinkage of the cells might be the reason behind the rapid increase in ERK phosphorylation [30]. Although we detected increased cell spreading in our system, we did not assess cell size immediately after AA starvation.

α2β1 integrin has been found to be upregulated in some types of cancer. Indeed, α2β1 integrin expression was increased in metastatic breast cancer cells compared to non-metastatic ones. α2 integrin overexpression in MDA-MB-231 cells enhanced their invasiveness and migration, but not their proliferation in vitro; however, it enhanced both their tumour growth and metastasis to the bone in vivo [45]. On the contrary, the loss of α2 in MMTV-neu mice increased tumour metastasis to the lungs in vivo, without affecting tumour growth [46]. Moreover, α2 integrin expression has been found to be upregulated in KRAS-mutated pancreatic cancer, compared to KRAS WT [40] and α2 integrin overexpression boosted the proliferation of PDAC cells in vitro and the tumour volume in vivo [47].

Here we showed that AA deprivation upregulated α2 integrin expression, in both 2D and 3D systems. This was mediated by the RAS-MEK-ERK pathway in cells harbouring RAS mutations and, in turn, increased cells adhesion, collagen I uptake and migration. Together, these data indicate that the nutrient-starved breast and pancreatic TME potentiates mutant RAS signalling, promotes nutrient scavenging and supports cell growth and migration, suggesting that α2 integrin could represent a promising therapeutic target for mutant RAS tumours.

## Methods

### Reagents

DMEM, high glucose, pyruvate; DMEM/F-12, glutamax; dialyzed foetal bovine serum (FBS), horse serum (HS) were from Gibco; DMEM, no amino acids from US Biological life sciences; FBS, Hydrocortisone and Insulin solution human from Sigma-Aldrich; Penicillin/ Streptomycin (Pen/strep) from Life technologies; Rat tail high concentration Collagen type I from Corning; Growth factor-reduced Matrigel from SLS; Hoechst 33342 from Invitrogen; NHS-Fluorescein, Phalloidin Alexa Fluor 555, Paraformaldehyde and pHrodo^TM^ iFL Red STP ester from ThermoFisher; Vectashield mounting reagent with DAPI from VECTOR laboratories; 4-15% Mini PROTEAN TGX stain Free Protein gels from BioRad Laboratories; Protein ladder “Color Prestained Protein Standard” from New England Biolabs; High-Capacity cDNA Reverse Transcription kit from Applied biosystems; qPCRBIO SyGreen Blue Mix Lo-ROX from PCR BIOSYSTEMS; FITC-Anti Human CD49b (ITGA2) Antibody from Biolegend (#359306); Mouse anti-human CD49b (ITGA2) antibody from BD biosciences (#611017); P44/42 MAPK (ERK1/2) antibody (#9102) and Phospho-p44/42 MAPK (ERK1/2) antibody (#9101) from Cell signalling; GAPDH antibody from Santa Cruz Biotechnology (#sc-47724); α-Tubulin was from Cell signalling (#3873); IR Dye 680LT anti-Rabbit antibody and IR Dye 800LT anti-Mouse antibody from LICOR Biosciences; E64d (Aloxistatin) from AdooQ Bioscience; BTT-3033 and Torin-1 from Tocris Biosciences; MRTX1133, GCN2iB and RMC-6236 from MedChemExpress; Selumetinib from Selleck Chemicals.

### Cell culture

MCF10CA1 (RRID:CVCL_6675, female) cells were cultured in high glucose Dulbecco’s Modified Eagle’s Medium (DMEM)/F12 medium supplemented with 5% horse serum (HS), 1% penicillin and streptomycin (Pen/strep), 0.2 μg/ml Hydrocortisone, 20 ng/ml EGF and 10 μg/ml Insulin. MDA-MB-231 (ATCC HTB- 26, RRID:CVCL_0062, female), MCF7 (RRID:CVCL_0031, female), PANC-1 (RRID:CVCL_0480, male) and SW1990 (RRID:CVCL_1723, male) cells were cultured in high glucose DMEM supplemented with 10% FBS and 1% Pen/strep. All cells were kept at 37°C and 5% CO_2_ and passaged twice a week. MCF10CA1 cells were kindly provided by Prof Giorgio Scita, IFOM, Milan (Italy). PANC-1 and SW1990 cells were kindly provided by Dr Helen Matthews (University of Sheffield). MCF7 cells were kindly provided by Prof Penelope Ottewell (University of Sheffield). All cells were regularly tested for mycoplasma. Starvation media compositions are reported in Table 1. MCF10CA1 cell starvation and complete media were supplemented with HS, while for all the other cell lines, starvation and complete media were supplemented with dialysed FBS (DFBS).

**Table 1.** Composition of starvation media used for each cell line

| Cell line | Starvation condition | Media composition |
| --- | --- | --- |
| MCF10CA1 | Amino acid starvation | (+) DMEM, (+) 4.5 g/L D-glucose, (+) 5% HS, (+) 1% Pen/Strep, (+) 20 ng/ml EGF, (+) 10 μg/ml insulin, (+) 0.2 μg/ml hydrocortisone |
| MDA-MB-231<br>MCF7<br>SW1990<br>PANC-1 | Amino acid starvation | (+) DMEM, (+) 4.5 g/L D-glucose, (+) 10% DFBS, (+) 1% pen/strep |
| SW1990 | Tumour Interstitial Fluid Medium (TIFM) | Kindly provided by Muir lab [3] (+) 10% DFBS, (+) 1% pen/strep |

### ECM preparation

A solution of 2 mg/ml collagen I or 3 mg/ml Matrigel in PBS was prepared, 15 μl/well were added to a 96-well plate and left to polymerise at 5% CO_2_, 37°C. PBS was added after 4 hours, and plates were kept in the incubator overnight. For 35mm glass-bottom dishes, a solution of 0.5-1 mg/ml collagen I in PBS was prepared, the dishes were coated with 100 μl/dish and kept at 37°C to polymerise. PBS was added after 1 hour, and dishes were kept in the incubator overnight.

### ECM uptake

35mm glass-bottom dishes were coated with collagen I. Collagen I was labelled with 20 μg/ml pH-rodo iFL red in PBS containing 0.1 M NaHCO_3_ and incubated for 1 hour at RT. Then, collagen I was washed twice with PBS. PBS was added and the dishes were left at 5% CO_2_, 37°C overnight. Where indicated, cells were incubated in starvation media overnight, in the presence or absence of 10 μM Selumetinib. The day after cells were seeded on collagen I-coated dishes and incubated at 5% CO_2_, 37°C for 6 hours. Where indicated, cells were incubated for 2 hours then 15 μM BTT-3033 was added to the cells for 4 hours. 1:1000 Hoechst 33342 was spiked into the media and incubated for 15 minutes. Media was changed with fresh media, and cells were imaged live with a Nikon A1 confocal microscope, 60x oil objective. Fiji/Image J software [48] was used to quantify ECM uptake as in [7, 20]. Briefly, the cell area was measured by drawing around the cells using the brightfield channel. The channel relative to the ECM staining then was thresholded to remove the background signal, ‘particle analysis’ was performed. The % area covered by collagen I fluorescence signal was plotted as ‘collagen I uptake index’.

### Cell proliferation

96-well plates were coated with collagen I. Cells were seeded in complete media and plates were incubated for 4 hours at 5% CO_2_, 37°C. Media was removed, cells were washed twice with PBS and media was changed to starvation media containing DMSO, 10 or 15 μM BTT-3033, where indicated. Inhibitor/DMSO were added every other day for 4 or 7 days. Cells were fixed with 4 % PFA for 15 minutes at RT, stained with 1:1000 Hoechst 33342 for 15 minutes and imaged with an ImagesXpress micro, 2x objective and DAPI filter in the Sheffield RNAi Screening Facility (SRSF). The images were obtained by acquiring multiple sites to cover the entire wells (4 sites/well for the 2x objective). The cell number was measured using MetaXpress and Costume Module Editor software (CME) by designing an algorithm to detect the nuclei by covering the range of nuclei size and intensity.

### 3D culture

5 μl of growth factor-reduced Matrigel was added to each well of 8-well glass-bottom chamber slides, spread evenly, and incubated at 5% CO_2_, 37°C for 15-30 minutes. Cells were trypsinised, counted and diluted in a solution of media containing 2% Matrigel. Cells were seeded on the Matrigel coated wells to a final concentration of 1.25x 10^4^ cells/ml. The day after, media was changed to complete or starvation media in the presence or absence of the indicated inhibitors and cells were incubated at 5% CO_2_, 37°C for 2 days. Cells were fixed with 4% PFA, permeabilised with 0.25% Triton X-100, washed twice with PBS then blocked with 1% Bovine Saline Albumin (BSA) for 1 hour at RT. Cells were washed twice with PBS and incubated for 1 hour at RT with 1:200 anti-human FITC-conjugated α2 integrin antibody in PBS. Cells were washed twice with PBS and two drops of Vectasheild containing DAPI were added. Chambers were sealed and imaged with a Nikon A1 confocal microscope, 60x oil objective. The mean fluorescence intensity was measured using Fiji/Image J.

### Western blotting

1x10 ^6^ Cells were seeded in 6-well plates. After a 4-hour incubation at 5% CO_2_, 37°C, cells were washed twice with PBS, media was changed to complete or starvation media in the presence of the indicated inhibitors and cells were incubated at 5% CO_2_, 37°C for 18 hours. Media was removed, cells were washed twice with ice-cold PBS and 100 μl/well ice-cold lysis buffer (50mM Tris and 1% SDS in water) was added. Cells were scraped; the lysate was transferred to a QiaShredder column and spun at full speed for 5 minutes at RT. The flowthrough was transferred to a clean Eppendorf tube. Loading buffer (1mM DTT in 4x NuPage) was added to the samples and heated at ∼70°C for 5 minutes. The samples were loaded in 4-15% gradient gels, which were run (25mM Tris, 192mM Glycine and 1% SDS in dH2O) at 100V for 1 hour 15 minutes. Proteins were transferred (10% 10X Towbin buffer, 20% methanol) to IMMOBILON-FL-PVDF membranes at 100V for 1 hour 15 minutes. The membranes were blocked for 1 hour with 5% BSA in TBS-T at RT then incubated with 1:1000 Mouse anti human CD49b (ITGA2) or GAPDH primary antibodies in 5% BSA in TBS-T overnight in a cold room. The day after, the membrane was washed 3 times with TBS-T, 10 minutes each, and incubated with IR Dye 680LT anti Rabbit IgG antibody and IR Dye 800LT anti Mouse IgG antibody Licor secondary antibodies in TBS-T+ 0.01% SDS for 1 hour at RT on a rocker. Membranes were washed 3 times with TBS-T, 10 minutes each, rinsed with water and imaged with a Licor Odyssey Sa imager. The signal intensity was measured using Image Studio Lite software.

To measure ERK phosphorylation, cells were seeded in 6-well plates. After a 4-hour incubation at 5% CO_2_, 37°C, cells were washed twice with PBS, media was changed to media without serum and growth factors and cells were incubated at 5% CO_2_, 37°C overnight. The day after, media was removed, cells were washed with complete or starvation media, without serum and growth factors and the corresponding media were added. Cells were lysed after 5, 10, or 15 minutes and samples were run as above. Alternatively, cells were seeded in 6-well plates. After a 4-hour incubation at 5% CO_2_, 37°C, cells were washed twice with PBS, media was changed with complete media without serum and growth factors and cells were incubated at 5% CO_2_, 37°C overnight. The day after 200 nM MRTX1133 or 10 μM Selumetinib were added for 30 minutes then media was changed to inhibitor-containing starvation media. Cells were lysed after 5 minutes and samples were run as above. The membranes were blocked for 1 hour with 5% BSA in TBS-T at RT then incubated with 1:1000 P44/42 MAPK (ERK1/2), Phospho- p44/42 MAPK (ERK1/2), GAPDH or α-Tubulin primary antibodies in 5% BSA in TBS-T overnight in a cold room. The day after, the membrane was washed 3 times with TBS-T, 10 minutes each, and incubated with the secondary antibodies for 1 hour at RT on a rocker. Membranes were washed 3 times with TBS- T, 10 minutes each, rinsed with water and imaged with a Licor Odyssey Sa imager. The signal intensity was measured using Image Studio Lite software.

### siRNA-mediated knockdown

For cell proliferation assays, 96-well plates were coated with collagen I. A mix of 5 μl serum-free media and 5 μl of 5 μM siRNA (Table 2) was added to each well and the plates were incubated for 5 minutes at RT. A mix of 0.2μl Dharmafect 1 (DF1) and 4.8 μl serum-free media was prepared and incubated for 5 minutes at RT. The DF1 mix was added to each well of 96-well plates (final concentration of siRNA is 30 nM) and incubated for 20 minutes at RT on a gentle rocker. 2x10^3^ cells in 80 μl media containing 5% horse serum were added and incubated at 5% CO_2_, 37°C overnight. The day after, the media was removed, cells were washed twice with PBS and media was changed to starvation media.

**Table 2.**
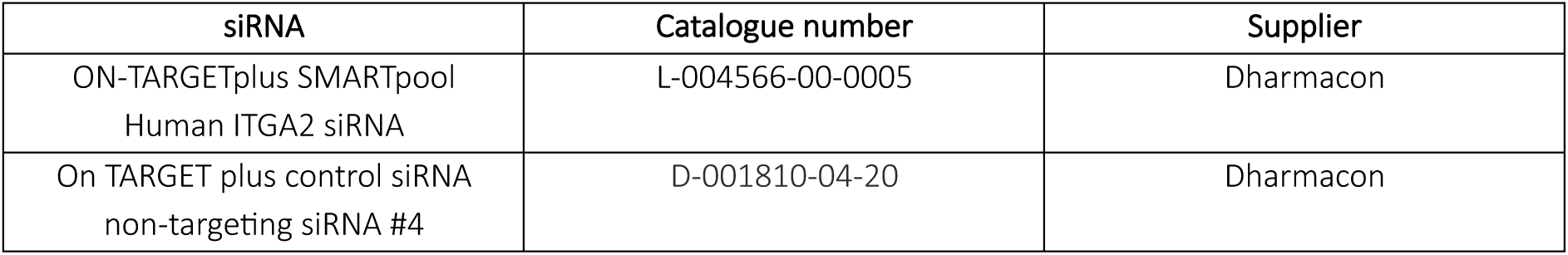
siRNAs used in this study.

| siRNA | Catalogue number | Supplier |
| --- | --- | --- |
| ON-TARGETplus SMARTpool<br>Human ITGA2 siRNA | L-004566-00-0005 | Dharmacon |
| On TARGET plus control siRNA<br>non-targeting siRNA #4 | D-001810-04-20 | Dharmacon |

Alternatively, a mix of 197 μl of serum free media and 3 μl of 5 μM siRNA was added to each well of 6- well plates and incubated for 5 minutes at RT. A mix of 4 μl of Dharmafect 1 (DF1) and 196 μl of serum-free media was prepared and incubated for 5 minutes at RT. The DF1 mix was added to each well of 6- well plate and incubated for 20 minutes at RT on a gentle rocker. 5x10^5^ cells in 1.6 ml of media containing 5% horse serum were added and the cells were incubated at 5% CO_2_, 37°C for 72hrs. Cells were collected, and analysed by Western Blotting.

### RT-qPCR

1x10^6^ cells were seeded in 6-well plates in complete media, after 4 hours of incubation at 5% CO_2_ and 37°C, media was changed to complete or starvation media in the presence or absence of 5μM GCN2iB, 200 nM MRTX1133 or 10 μM Selumetinib and incubated overnight. The day after, media was removed, cells were washed twice with PBS and trypsinised. Cells were collected and centrifuged at 1000 rpm for 5 minutes. The supernatant was removed, and the cell pellets were resuspended with 1 ml PBS and centrifuged at 1400rpm for 5 minutes. The PBS was removed, and the pellets were kept at-70°C. Using QIAGEN RNA extraction kit, the mRNA was extracted according to the manufacturers’ protocol. The mRNA concentration was measured by a Nanodrop LITE Spectrophotometer (Thermo Scientific), and the extracted mRNA was used to synthesise cDNA. The following master mix was prepared by using High-Capacity cDNA Reverse Transcription kit (Table 3): 5.8 μl master mix per sample was added to a 14.2 μl mix of nuclease free water + 1000 ng of RNA (-RT control was prepared using the sample with the highest RNA concentration). The samples were run in a thermal cycler, and the synthesised cDNA was kept at -70°C. The cDNA was then diluted 1:10 in nuclease free water and 3 μl of the cDNA solution was added to the master mix, which was prepared using qPCRBIO SyGreen Blue Mix (Table 4).

**Table 3.** RT master mix composition.

| Component | Volume per sample |
| --- | --- |
| 10x RT Random Primers | 2.0 µl |
| 25x dNTP Mix | 0.8µl |
| 10x RT buffer | 2.0µl |
| MultiScribe™ Reverse Transcriptase (RT) | 1.0µl |

**Table 4.** QPCR master mix composition.

| Component | Volume | Final concentration |
| --- | --- | --- |
| 2x SYBR Green PCR Master Mix | 5µl | 1x |
| 10x QuantiTect Primer Assay (table 5) | 1µl | 1x |
| Water | 1µl |  |

**Table 5.** List of primers

| Primer | GeneGlobe Id | Catalogue number | Supplier |
| --- | --- | --- | --- |
| Hs_ITGA2_1_SG QuantiTect Primer Assay | QT00086695 | 249900 | Qiagen |
| Hs_ITGA3_1_SG QuantiTect Primer Assay | QT00047404 | 249900 | Qiagen |
| Hs_ITGA5_1_SG QuantiTect Primer Assay | QT01008126 | 249900 | Qiagen |
| Hs_ITGA6_1_SG QuantiTect<br>Primer Assay | QT00025725 | 249900 | Qiagen |
| Hs_ITGB1_1_SG QuantiTect<br>Primer Assay | QT00068124 | 249900 | Qiagen |
| Hs_GAPDH_1_SG QuantiTect<br>Primer Assay | QT00079247 | 249900 | Qiagen |

The samples were loaded in 384-well plates (3 technical replicates per sample) including the-RT control and a water control for each tested gene. The plate was sealed and spun down for few seconds. The samples then were run using a QuantStudio 12K Flex Real-Time PCR System and the results were obtained using the following equation:

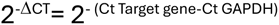

### Cell adhesion

96-well plates were coated with 2 mg/ml collagen I or 3 mg/ml Matrigel in PBS. Cells were starved in AA-free media or kept in complete media for 18 hours in the presence or absence of 200 nM MRTX1133 or 10 μM Selumetinib. Cells were seeded on ECM or plastic for 15, 30, or 60 minutes. Cells were washed twice with PBS, fixed with 4% PFA and stained with 1:1000 Hoechst 33342 for 15 minutes at RT. Cells were washed twice with PBS and imaged with an ImageXpress micro, 2x objective, or an INCell Analyser 2200. Cell number was measured using MetaXpress and CME software or with Cell Profiler. Alternatively, 12 well plated were coated with 1mg/ml collagen 1 in PBS. Cells were starved in AA-free media or kept in complete media for 18 hours and seeded on collagen I. Cells were imaged live with a 10x Nikon Inverted Ti eclipse with Oko-lab environmental control chamber (10-minute intervals) and analysed using Fiji/Image J.

### Wound healing assay

Cells were detached using TrypLe express and neutralised using 10% FBS media. Cells (1/10 of a confluent 10cm dish) were seeded into 12-well plates and incubated overnight to achieve a monolayer. The next morning, a 0.5mg/ml collagen I solution in complete or starvation media was prepared for the overlay. Monolayers were washed once using either complete or starvation media. Cells were then scratched manually by creating a cross using the end of a 200 μl pipette tip and washed again two times with corresponding complete or starvation media. Collagen I mixture was then laid over the scratched monolayers and left in the incubator for 30 minutes to polymerise, after which wells were topped up with complete or starvation media. Each branch of the scratches was imaged at 0h, 2h, 4h, 6h and 8h using an Olympus Inverted Fluorescence Microscope, 4x objective. Scratch areas were manually measured by tracing the perimeter along the invasive front using Fiji/Image J. Data were then normalised dividing the remaining gap area over the initial scratch area.

### Survival analysis

The survival analysis was performed by GEPIA2 (Gene Expression Profiling Interactive Analysis, RRID:SCR_018294 http://gepia2.cancer-pku.cn/#survival) using RNA sequencing expression data of 9,736 tumours and 8,587 normal samples from the TCGA and the GTEx projects, and a standard processing pipeline [49].

### Statistical analysis

Graphs were created in GraphPad prism 10 software. Mann-Whitney test was used to compare between two groups with one variable. One-way ANOVA (Kruskal-Wallis, Dunn’s multiple comparisons test) was applied to compare between three or more groups with one variable. Two-way ANOVA (Tukey’s multiple comparisons test) was applied to compare between groups with two independent variables.

## Acknowledgements

Imaging work was performed at the Wolfson Light Microscopy Facility, University of Sheffield (funded by the Wellcome Trust, grant WT093134AIA), using a Nikon A1 confocal microscope and a Nikon Inverted Ti eclipse with Oko-lab environmental control chamber. High-throughput imaging was performed in the RNAi facility at the University of Sheffield. The QPCR machine was part of the biomedical research equipment at the University of Sheffield.

## Competing interests

The authors have declared that no competing interests exist.

### Authors’ contributions

Conceptualisation: ER, BY Data curation: BY Formal analysis: BY, MN, ZB, RB, IO Funding acquisition: ER Investigation: BY, MN, ZB, RB, IO Methodology: BY, RB

Project administration: ER Supervision: ER

Writing, review and editing: BY, ER

---

**Figure S1.**
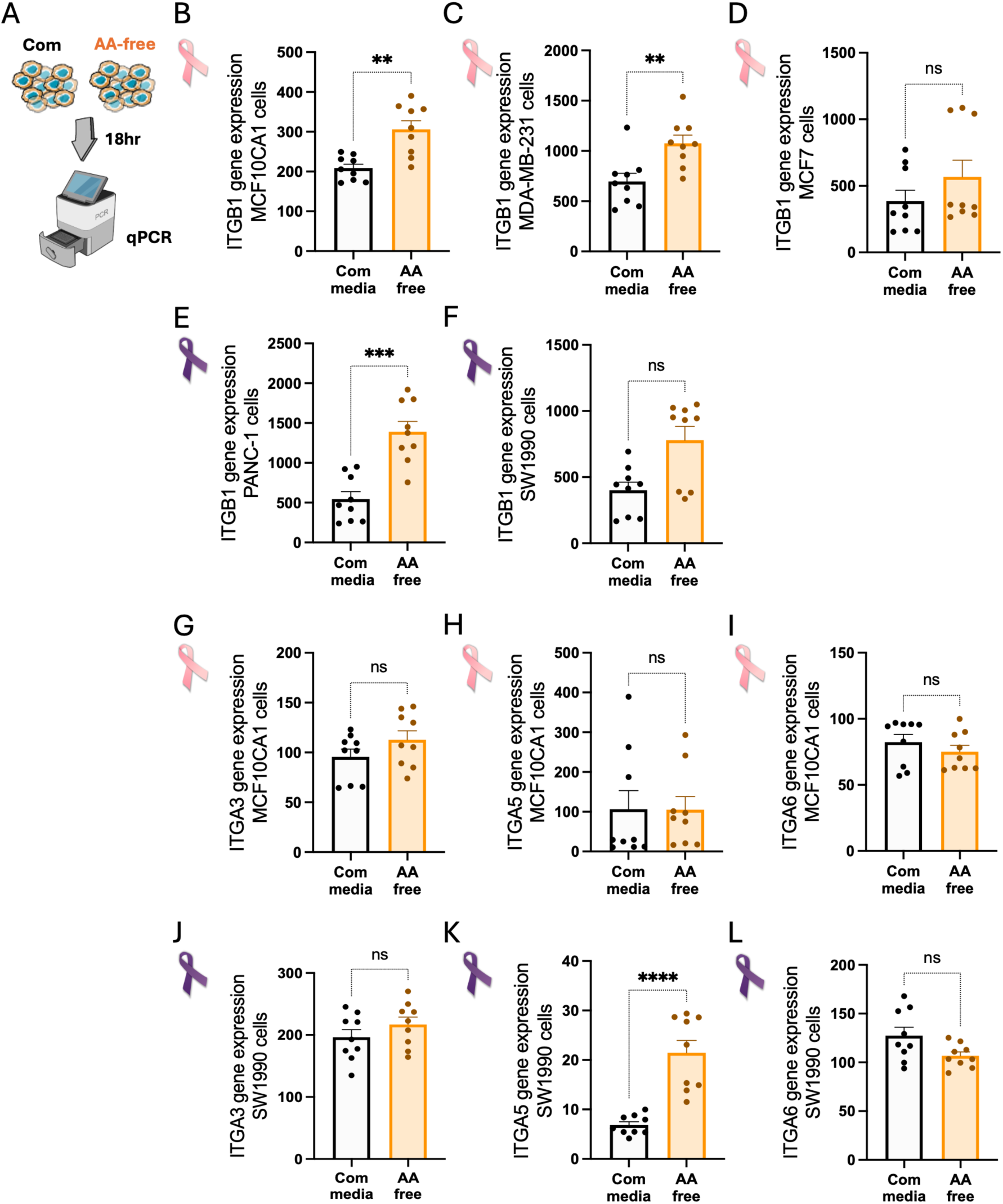
Amino acid starvation promoted. β**1 integrin expression in a cell line-dependent manner, without affecting**α**3,** α**5 or** α**6. (A)** Schematic representation of experimental set up. Breast cancer cells **(B,G-I)** MCF10CA1, **(C)** MDA-MB-231 and **(D**) MCF7 and pancreatic cancer cells **(E)** PANC-1 and **(F,J-L)** SW1990 were seeded on plastic in complete media (Com) or amino acid-free media (AA) for 18 hours. mRNA was extracted and β1 integrin (ITGB1, B-F), α3 integrin (ITGA3, G,J), α5 integrin (ITGA5, H,K), and α6 integrin (ITGA6, I,L) expression was measured by SYBR-Green qPCR. GAPDH was used as a control, and the data were plotted as 2^-ΔCT^. Graphs show mean ± SEM. N=3 independent experiments, ns: non- significant, **p < 0.01, ***p < 0.001, ****p < 0.0001 Mann-Whitney test. Pink and purple ribbons are from wikicommons, http://creativecommons.org/licenses/by-sa/3.0/. Illustrations are with items from NIAID NIH BIOART, bioart.niaid.nih.gov/bioart/426.

**Figure S2.**
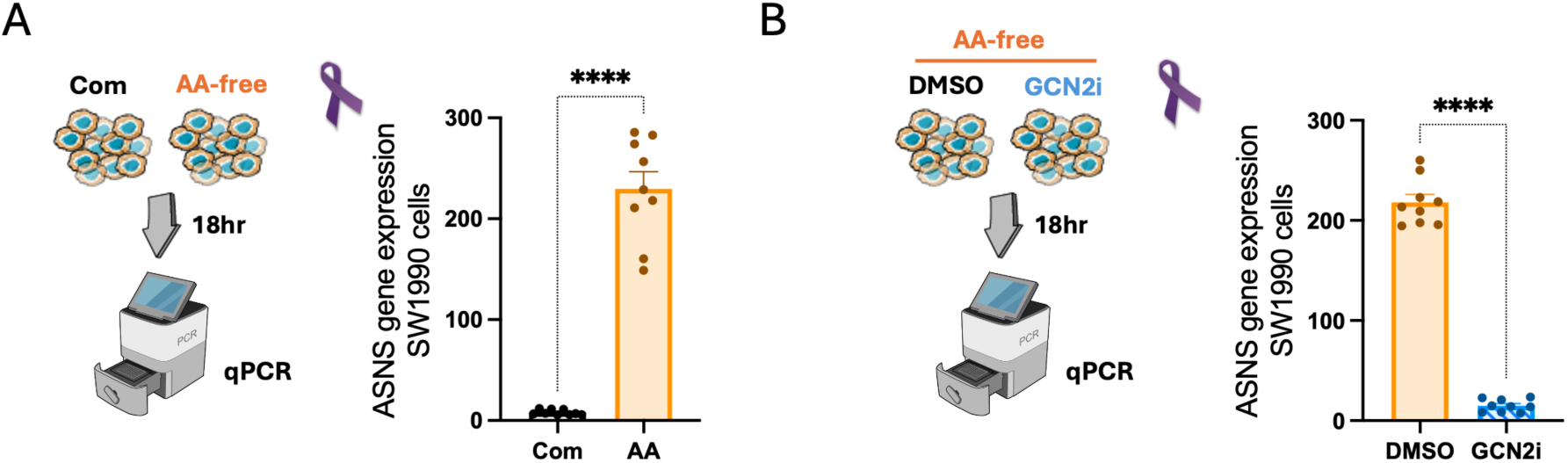
GCN2 inhibition completely abolished amino acid starvation-induced expression of ASNS. (A) SW1990 cells were seeded in complete media (Com) or amino acid-free media (AA) for 18 hours. (B) SW1990 cells were seeded in amino acid-free media (AA) for 18 hours in the presence of 5 μM GCN2iB (GCN2i) or DMSO as a control. mRNA was extracted and ASNS expression was measured by SYBR-Green qPCR. GAPDH was used as a control and the data were plotted as 2-^ΔCT.^ N=3 independent experiments, values represent the Mean ± SEM. ****p < 0.0001 Mann-Whitney test. Purple ribbons are from wikicommons, http://creativecommons.org/licenses/by-sa/3.0/. Illustrations are with items from NIAID NIH BIOART, bioart.niaid.nih.gov/bioart/426.

**Figure S3.**
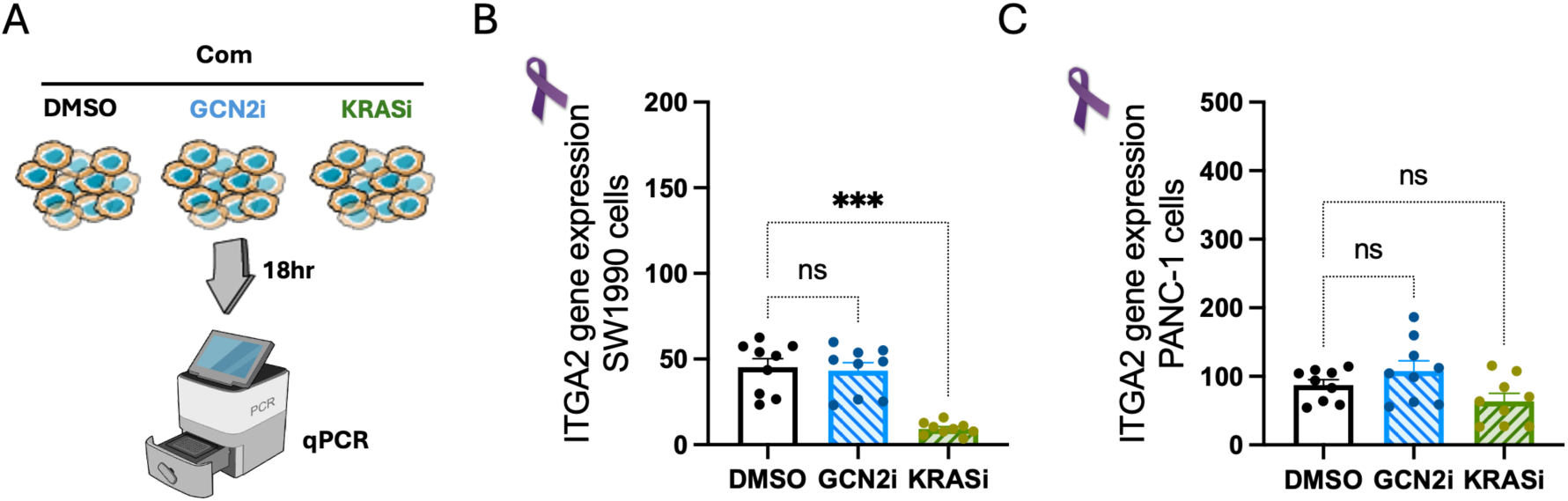
KRAS, but not GCN2, controlled. α**2 integrin expression in complete media in a cell line-dependent manner. (A)** Schematic representing the experimental approach. **(B, C)** Pancreatic cancer cells (B) SW1990 and (C) PANC-1 were seeded in complete media for 18 hours in the presence of 5 μM GCN2iB (GCN2i), 200 nM MRTX1133 (KRASi) or DMSO as a control. mRNA was extracted and α2 integrin (ITGA2) expression was measured by SYBR-Green qPCR. GAPDH was used as a control, and the data were plotted as 2^-ΔCT^. Graphs show mean ± SEM. N=3 independent experiments, ns: non-significant, ***p = 0.0002 Kruskal Wallis, Dunn’s multiple comparisons test. Pink and purple ribbons are from wikicommons, http://creativecommons.org/licenses/by-sa/3.0/. Illustrations are with items from NIAID NIH BIOART, bioart.niaid.nih.gov/bioart/426.

**Figure S4.**
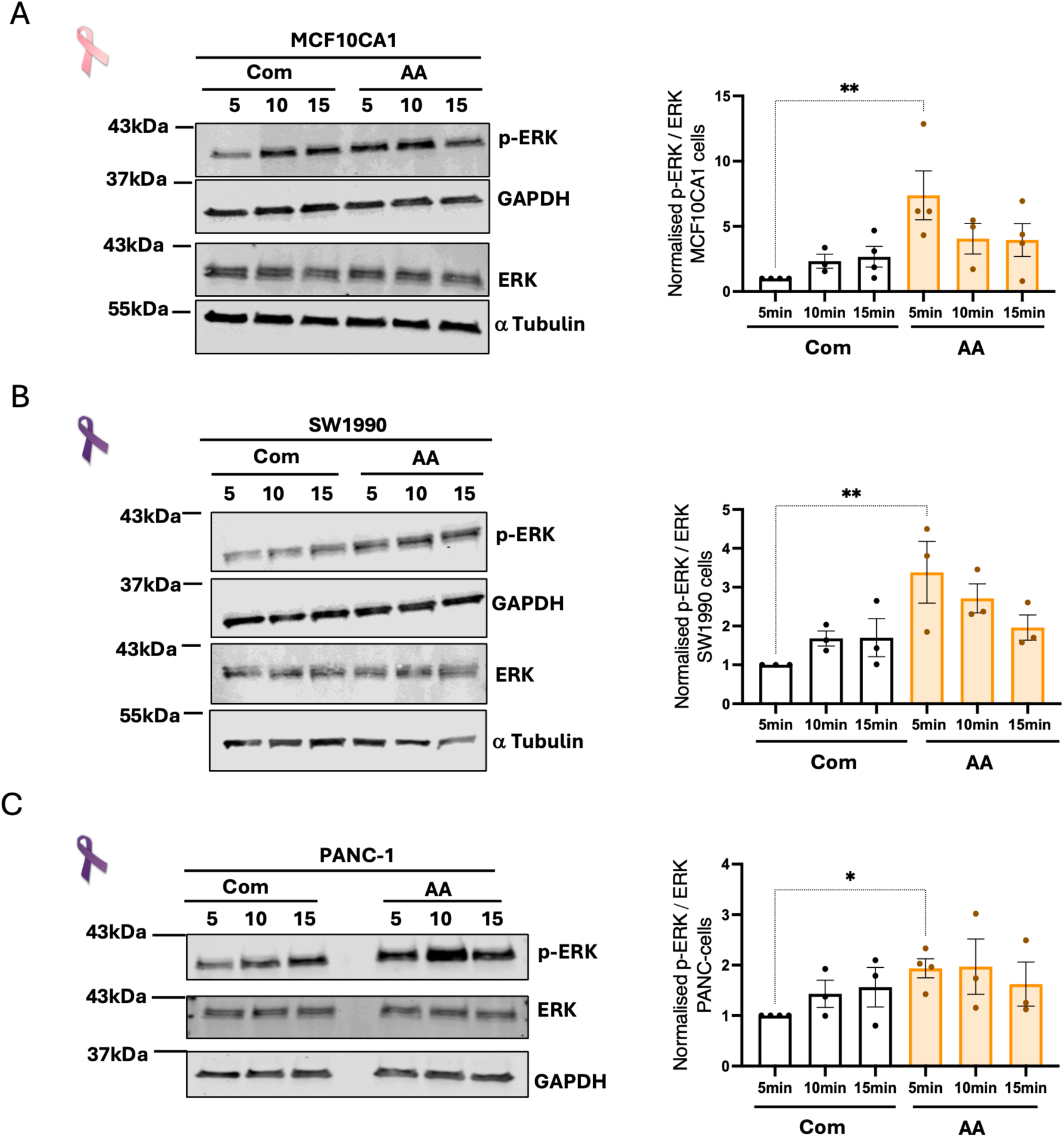
Amino acid starvation promoted ERK phosphorylation. **(A)** MCF10CA1 cells, **(B)** SW1990 cells and **(C)** PANC-1 cells were seeded in complete media, the media was changed after 4 hours to serum-free media and cells were incubated for 18 hours. Media was changed to serum-free complete or serum- and A- free media, cell lysates were collected after 5, 10, or 15 minutes, and the samples were run by Western Blotting. Membranes were stained for ERK, phospho-ERK (p-ERK), α-Tubulin and GAPDH and were imaged with a Licor Odyssey Sa system. The protein band intensity was measured with Image Studio Lite software and the p-ERK/ERK ratio was plotted, normalised to 5 min complete media. N≥3 independent experiments, graphs show mean ± SEM. *p < 0.05, ** p < 0.01 Kruskal-Wallis, Dunn’s multiple comparisons test. Pink and purple ribbons are from wikicommons, http://creativecommons.org/licenses/by-sa/3.0/. Illustration is with items from NIAID NIH BIOART, bioart.niaid.nih.gov/bioart/580.

**Figure S5.**
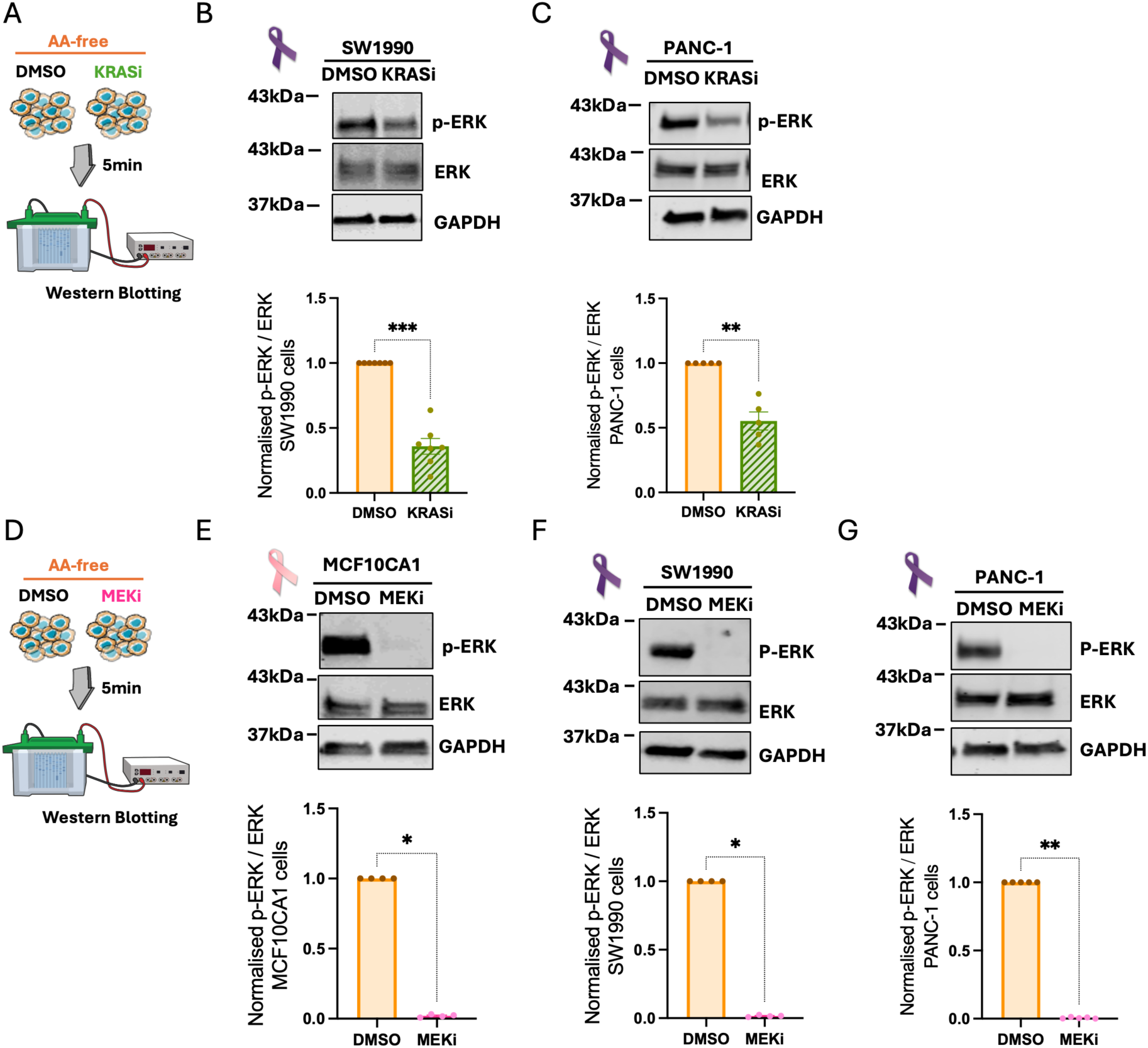
KRAS and MEK pharmacological inhibition blunted ERK phosphorylation. (A, **D)** Schematic representation of experimental set up. **(B, F)** SW1990, **(C, G)** PANC-1 and **(E)** MCF10CA1 cells were seeded in complete media, the media was changed after 4 hours to serum-free complete media and cells were incubated for 18 hours. 10 μM Selumetinib (MEKi), 200 nM MRTX1133 (KRASi) or DMSO as a control were added for 30 minutes. Media was changed to serum- and AA-free media with the inhibitors. The cell lysates were collected after 5 minutes, and the samples were run by Western Blotting. Membranes were stained for ERK, phospho-ERK (p-ERK), and GAPDH and were imaged with a Licor Odyssey Sa system. The protein band intensity was measured with Image Studio Lite software and the ratio of p-ERK/ERK was plotted. N≥4 independent experiments, graphs show mean ± SEM. *p < 0.05, **p < 0.01 Mann Whitney test. Pink and purple ribbons are from wikicommons, http://creativecommons.org/licenses/by-sa/3.0/. Illustrations are with items from NIAID NIH BIOART, bioart.niaid.nih.gov/bioart/580.

**Figure S6.**
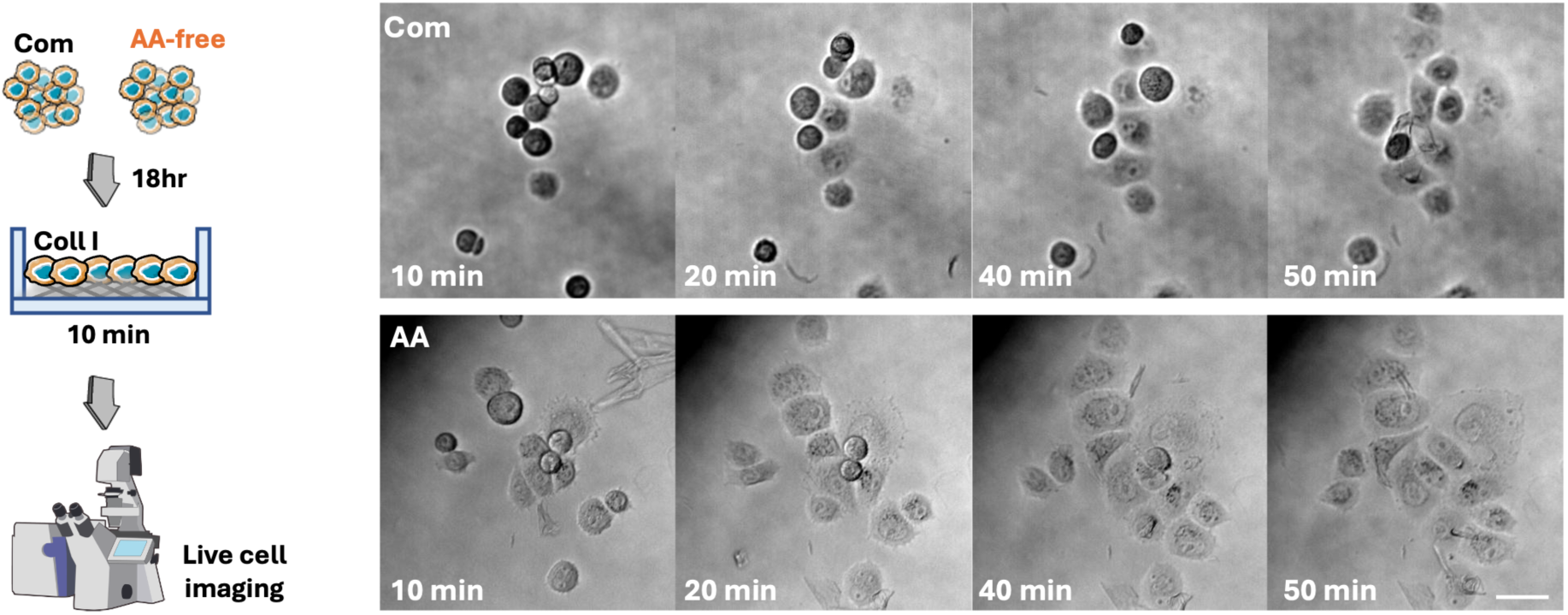
Amino acid starvation promoted PANC-1 cell spreading on collagen. **I.** PANC-1 cells were grown in complete media (Com, supplementary video 1) or amino acid free media (AA, supplementary video 2) for 18 hours. Cells were seeded on 1 mg/ml collagen I and imaged live with a 10x Nikon Inverted Ti eclipse with Oko-lab environmental control chamber (10-minute intervals). Representative still images extracted from the videos are shown. Scale bar 20 μm. Cell spreading area was measured with Fiji/Image J, representative of n=2 independent experiments. **p = 0.0014, ****p < 0.0001 Two-WAY ANOVA, Šídák’s multiple comparisons test. Illustrations in (A) are with items from NIAID NIH BIOART Source, bioart.niaid.nih.gov/bioart/86.

